# *Tenuivirus* uses a molecular bridge strategy to overcome insect midgut barriers for virus persistent transmission

**DOI:** 10.1101/404657

**Authors:** Gang Lu, Shuo Li, Changwei Zhou, Xin Qian, Qing Xiang, Tongqing Yang, Jianxiang Wu, Xueping Zhou, Yijun Zhou, Xin Shun Ding, Xiaorong Tao

## Abstract

Many persistent transmitted plant viruses, including *Rice stripe tenuivirus* (RSV), cause serious damages to crop productions in China and worldwide. Although many reports have indicated that successful insect-mediated virus transmission depends on proper virus–insect vector interactions, the mechanism(s) controlling interactions between viruses and insect vectors for virus persistent transmission remained poorly understood. In this study, we used RSV and its small brown planthopper (SBPH) vector as a working model to elucidate the molecular mechanism controlling RSV virion entrance into SBPH midgut for persistent transmission. We have now demonstrated that this non-enveloped *Tenuivirus* uses its non-structural glycoprotein NSvc2 as a helper component to bridge the specific interaction between virion and SBPH midgut cells, leading to overcome SBPH midgut barriers for virus persistent transmission. In the absence of this glycoprotein, purified RSV virion is not capable of entering SBPH midgut cells. In RSV-infected cells, glycoprotein NSvc2 is processed into two mature proteins: an amino-terminal protein NSvc2-N and a carboxyl-terminal protein NSvc2-C. We determined that NSvc2-N interacted with RSV virion and bound directly to midgut lumen surface via its N-glycosylation sites. Upon recognition by midgut cells, the midgut cells underwent endocytosis followed by compartmentalizing RSV virion and NSvc2 into early and then late endosomes. The acidic condition inside the late endosome triggered conformation change of NSvc2-C and caused cell membrane fusion via its highly conserved fusion loop motifs, leading to the release of RSV virion from endosome into cytosol. In summary, our results showed for the first time that a rice *Tenuivirus* uses a molecular bridge strategy to ensure proper interactions between virus and insect midgut for successful persistent transmission.

**Author summary:** Over 75% of the known plant viruses are insect transmitted. Understanding how plant viruses interacted with their insect vectors during virus transmission is one of the key steps to manage virus diseases worldwide. Both the direct and indirect virus–insect vector interaction models have been proposed for virus non-persistent and semi-persistent transmission. However, the indirect virus–vector interaction mechanism during virus persistent transmission has not been reported previously. In this study, we developed a new reverse genetics technology and demonstrated that the circulative and propagative transmitted *Rice stripe tenuivirus* utilizes a glycoprotein NSvc2 as a helper component to ensure a specific interaction between *Tenuivirus* virion and midgut cells of small brown planthopper (SBPH), leading to conquering the midgut barrier of SBPH. This is the first report of a helper component mediated-molecular bridge mechanism for virus persistent transmission. These new findings and our new model on persistent transmission expand our understanding of molecular mechanism(s) controlling virus–insect vector interactions during virus transmission in nature.

## Introduction

Arthropod insects play critical roles in epidemics of numerous animal and plant viruses [1–3]. Based on the mode of transmission, plant viruses can be classified into non-persistent, semi-persistent or persistent transmitted viruses [4–6]. For non-persistent and semi-persistent transmissions, plant viruses retain only inside insect stylets or on foregut surface for a short period of time. Upon probing or feeding on a new host plant, viruses are quickly injected into plant cells, together with insect saliva [7–9]. The persistent transmitted plant viruses (non-propagative or propagative) are required to enter insect vector bodies, and then circulate and/or replicate inside the vectors for several days to weeks. These persistent transmitted viruses need to pass insect midgut barrier, dissemination barrier, and then salivary gland barrier prior to be transmitted to new host plants [6, 10, 11]. Midgut is often considered to be one of major barriers for successful persistent transmissions of plant viruses. During the process of passing through barriers inside vectors, proper interactions between viruses and vectors are needed for successful transmission. However, the mechanism controlling the interactions between persistent transmitted plant viruses and their insect vector midgut barriers remains poorly understood.

Molecular bridge mechanism has been reported for non-persistent and semi-persistent transmitted plant viruses, respectively [4, 12, 13]. For example, virion of non-persistent or semi-persistent transmitted plant viruses were reported to interact with the cuticular proteins in the mouthparts or the foreguts [14, 15], and these virus–insect interactions required virally encoded non-structural helper factors as molecular bridges [12, 13]. Viruses in the genus *Potyvirus* are known to encode a helper component proteinase (HC-Pro) and this HC-Pro protein acts as a molecular bridge for potyvirus virion–aphid vector interactions [16–18]. Members in the genus *Caulimovirus* encode a different helper factor that helps virion to retain on insect maxillary stylet [19–21]. Although virion of multiple persistent transmitted plant viruses [e.g., *Luteovirus* [22, 23], *Geminivirus* [24, 25], *Reovirus* [26, 27], *Tospovirus* [28, 29], and plant *Rhabdovirus* [30, 31]] were also reported to bind directly to insect midgut cells, these bindings all depended on virion surface-exposed proteins. To date, no persistent transmitted (non-propagative or propagative) plants viruses has been reported for the requirement of additional helper proteins for the transmission.

*Rice stripe tenuivirus* (RSV) is known to be transmitted by small brown planthopper (SBPH) in a circulative and propagative manner, and causes severe rice losses in China and many other countries in Asia [32, 33]. The genome sequence of plant infecting *Tenuivirus* is similar to members of animal infecting *Phlebovirus* in the order of *Bunyavirales.* Most of the members in the order of *Bunyavirales* produce membrane-enveloped spherical virion with two surface-exposed glycoproteins. These surface-exposed glycoproteins are key determinants for entering host cells or for vector transmission [29, 34, 35]. However, virion of tenuiviruses are filamentous and do not have enveloped membranes. Purified tenuivirus virion was previously reported to be non-transmissible by vector insects [36–38]. RSV also encodes a glycoprotein NSvc2, which can be further processed into an amino-terminal part protein known as NSvc2-N and a carboxyl-terminal part protein known as NSvc2-C [39, 40]. The RSV encoded glycoprotein NSvc2 was not found in the purified virion samples [41, 42]. Based on the above published reports we hypothesized that *Rice stripe tenuivirus* must use a quite different mechanism to overcome the midgut barriers for its insect transmission.

To validate this hypothesis, we conducted various experiments using RSV and SBPH as our working model. We have now determined for the first time that the rice *Tenuivirus* uses a unique molecular bridge strategy to overcome insect midgut barrier for virus persistent transmission. We have shown that in the absence of RSV non-structural glycoprotein NSvc2, RSV virion was unable to enter SBPH midgut cells. We found that this RSV non-structural glycoprotein NSvc2 acts as a critical molecular bridge to mediate the interaction between RSV virion and SBPH midgut. NSvc2-N, a processed product from NSvc2, interacts with RSV virion and binds directly to the midgut barrier. Upon the successful interaction, midgut cells undergo endocytosis followed by compartmentalization of RSV virion and NSvc2 complexes in early and then late endosomes. NSvc2-C, another processed product from NSvc2, triggers membrane fusion under the acidic condition inside the late endosomes to release RSV virion into cytosol. These new findings expand our understanding of interactions between virion and insect vectors during transmissions of plant and animal viruses.

## Results

### Association of NSvc2 protein with RSV virion in midgut of SBPH

To examine whether RSV NSvc2 plays role(s) in circulative RSV transmission, we first conducted a time course study on the co-localization of NSvc2 and RSV virion in the midgut of SBPH during acquisition. SBPHs were fed on RSV-infected rice seedlings and then collected at 4, 8, 16 and 24 h after feeding (30 SBPHs at each time point), respectively. The collected insects were dissected and analyzed for the presence of NSvc2 and RSV virion by double-immunolabeling methods using an antibody against the amino-terminal NSvc2 (NSvc2-N) or RSV virion-surface nucleocapsid protein (NP). As shown in Fig 1A that, after 4 h feeding on the RSV-infected rice seedlings, numerous RSV virion (green) had accumulated in the midgut lumen. In the same tissues, NSvc2 was also detected (red) and found to co-localize with RSV virion on the actin-labelled intestinal microvillus (blue) (Fig 1A). The overlapped coefficient (OC) value for the red and green labeling signal was 0.76 ± 0.03 at 4 h post feeding (Fig 1E), indicating that NSvc2 and RSV virion were localized close to each other. At 8 h post feeding, NSvc2 was found to co-localize with RSV virion in various sized vesicles-like structures in epithelial cells (Fig 1B). Analysis of OC value showed again that NSvc2 and RSV were indeed localized close to each other (Fig 1F). At 16 h post feeding, RSV virion was detected together with NSvc2 in cytoplasm of midgut epithelial cells, with an OC value of 0.73 ± 0.06 (Fig 1C and 1G). Even at 24 h post feeding, NSvc2 was still associated with RSV virion, with an OC value of 0.76 ± 0.04, and the virus had spread into the surrounding midgut epithelial cells (Fig 1D and 1H). These data showed that RSV-encoded glycoprotein NSvc2 was associated with RSV virion in SBPH midgut during insect feeding on RSV-infected rice plants.

**Fig 1.**
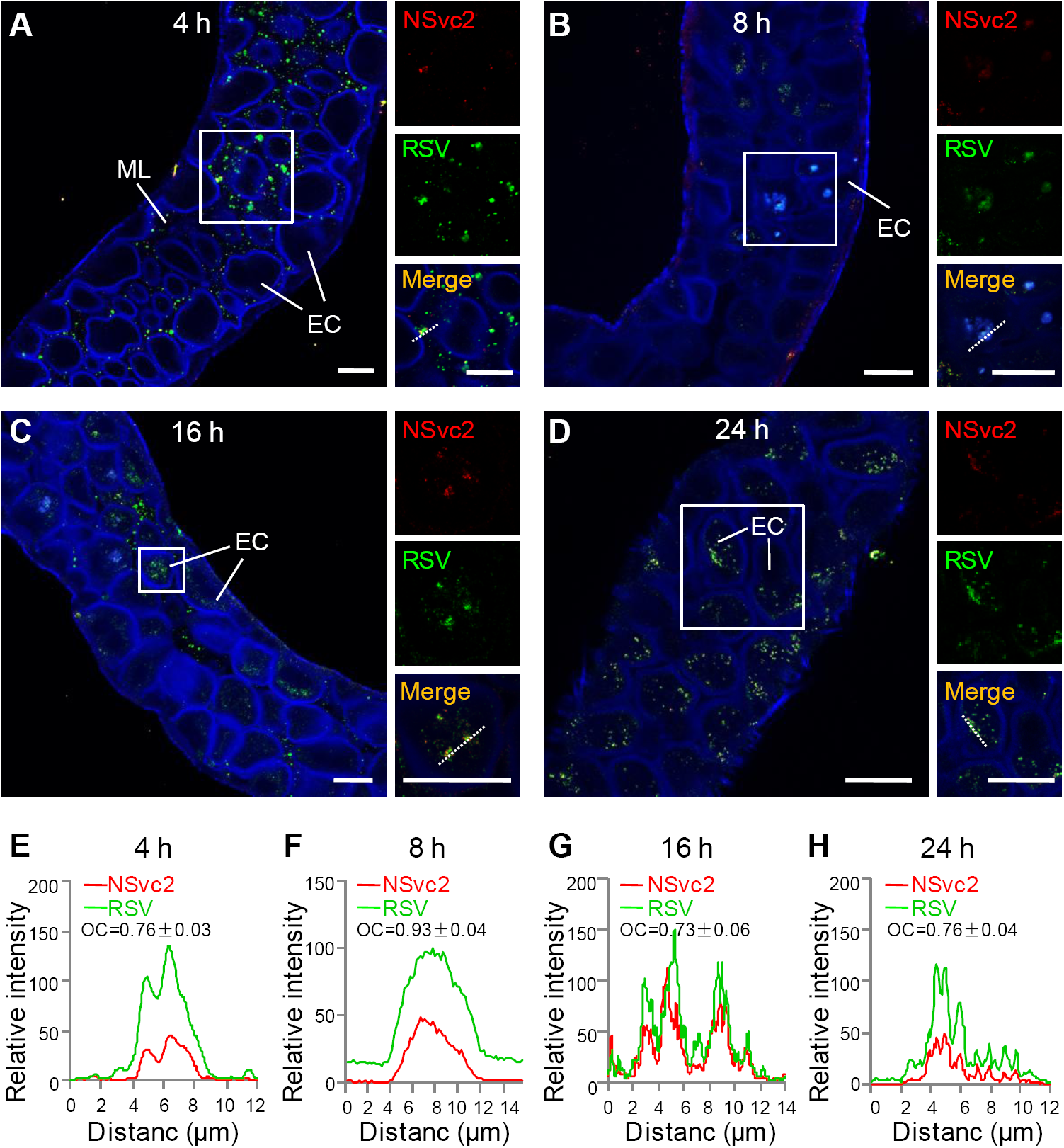
NSvc2 associates with RSV virion in SBPH midgut. (A) NSvc2 (Red) and RSV virion (green) were co-localized in midgut lumen and on the surface of intestinal microvillus (blue) at 4 h post feeding on RSV-infected rice plants. The presence of RSV virion, NSvc2, and actin were detected with antibodies specific for RSV NP, NSvc2-N, or actin. The boxed regions are shown on the right side of the corresponding images with three panels. The detection signal for NSvc2 is in red, the detection single for virion is in green and the merged detection signal is in yellow. (B) Co-localization of NSvc2 and RSV virion in vesicle-like structures in epithelial cells at 8 h post feeding. (C) Co-localization of NSvc2 and RSV virion in cytoplasm of epithelial cells at 16 h post feeding. (D) NSvc2 and RSV virion complexes were detected in epithelial cells of midgut at 24 h post feeding. (E–H) Analyses of overlapped fluorescence spectra from NSvc2 (red) and RSV virion (green) at different stages. Fluorescence signals were from the white dashed line indicated areas. The overlap coefficient (OC) values were determined individually using the LAS X software. ML, midgut lumen; EC, epithelial cell; Bar, 25 μm.

### NSvc2 protein is critical for RSV virion entrance into SBPH midgut

RSV virion was purified from virus-infected rice seedlings through ultracentrifugation using a 20% glycerol cushion. After ultracentrifugation, four different fractions starting from the top of the upper supernatant phase (Sup1 to Sup4), four fractions starting from the top of the lower 20 % glycerol phase (Gly1 to Gly4), and the pellet (Pel) were collected and analyzed individually by immunoblotting assays using an antibody against RSV NP or NSvc2-N (Fig 2A and 2B). Results showed that the pellet sample contained RSV virion, and the four supernatant fractions (Sup1 to Sup4) contained NSvc2 protein. In contrast, the four glycerol fractions (Gly1 to Gly4) contained RSV virion and NSvc2 protein (Fig 2B). Transmission Electron Microscopy showed that numerous filamentous RSV virion were present in the pellet sample (Fig 2C).

**Fig 2.**
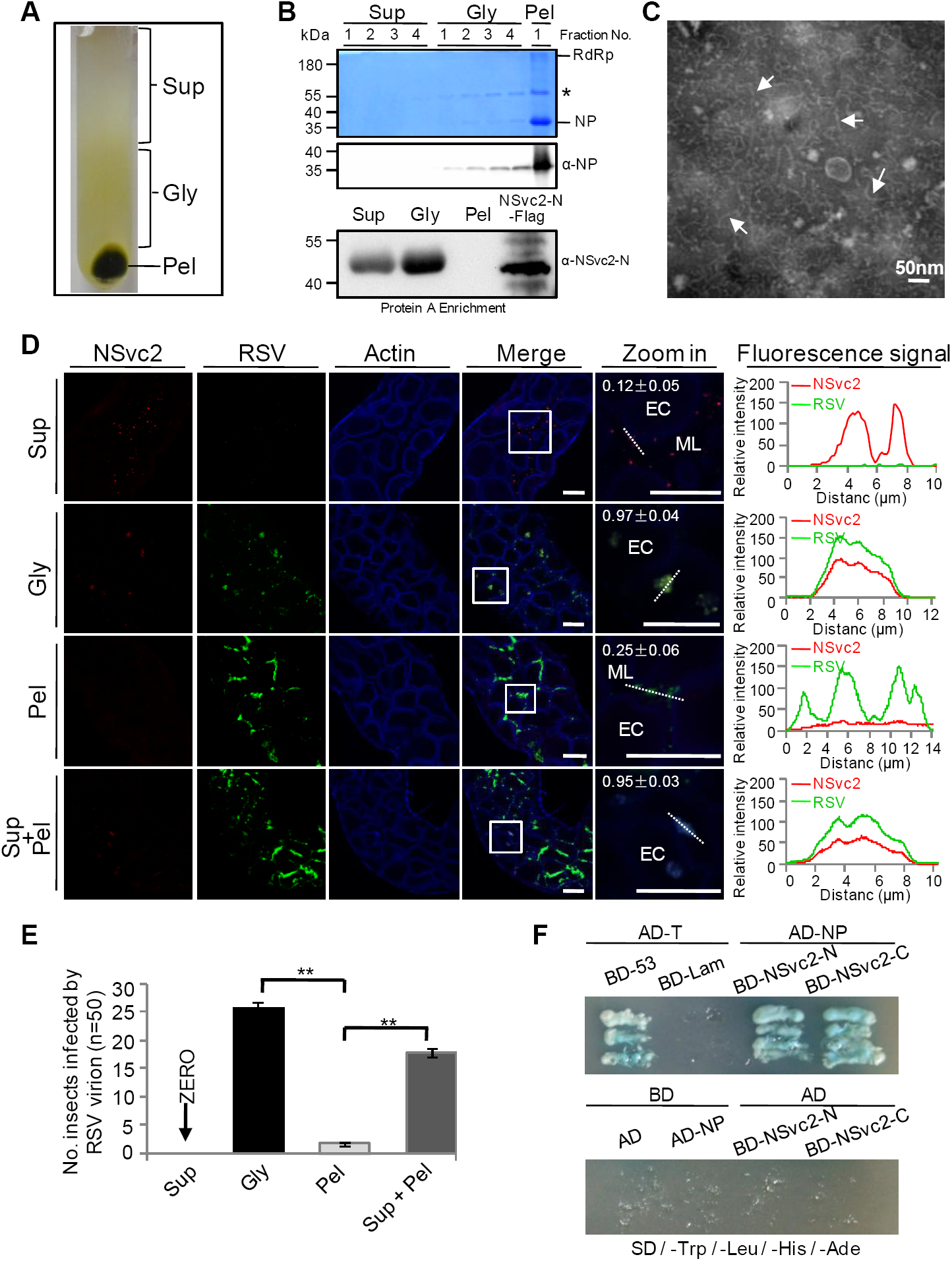
NSvc2 protein is required for RSV entrance into SBPH midgut. (A) NSvc2 is absent in purified RSV virion. Extract from RSV-infected rice plants was loaded on the top of 4 mL 20 % glycerol. The supernatant fractions, glycerol fractions and the pellet were collected after ultra-centrifugation. (B) The fractions were analyzed in gels followed by Coomassie blue staining or by immunoblotting using antibodies specific for RSV NP or NSvc2-N. Sizes of the protein bands are shown on the left. Asterisk indicates a RSV NP dimer. (C) Morphology of RSV virion in a resuspended pellet sample examined by Electron Microscopy. Arrows indicate filamentous RSV virion. Bar, 50 nm. (D) Immunofluorescence labeling of NSvc2 (red) and RSV virion (green) in SBPH midguts after feeding with the combined supernatant fraction, glycerol fraction, resuspended pellet or the supernatant fraction and resuspended pellet mixture. The boxed regions are enlarged and shown on the right side of the merged images. Overlapping fluorescence spectra analyses were done for the white dashed line indicated areas shown in the right panels. The overlap coefficient (OC) values were determined using the LAS X software. ML, midgut lumen; EC, epithelial cell; Bar, 25 μm. (E) Statistic analysis of RSV infection in SBPH after feeding on different fractions. Each bar represents three independent biological repeats from each experiment (n=50 / group). **, *p* < 0.01 by student *t*-test analysis. (F) Yeast two-hybrid assay for the interactions between RSV NP and NSvc2-N, or between RSV NP and NSvc2-C. RSV NP was fused to a GAL4 activation domain (AD-NP), and NSvc2-N or NSvc2-C was fused to a GAL4 binding domain (BD-NSvc2-N, BD-NSvc2-C). Yeast cells were co-transformed with indicated plasmids, and were assayed for protein interactions on synthetic dextrose-Trp/-Leu/-His/-Ade medium.

To verify the above findings, SBPHs were fed on a mixture of sucrose and the combined supernatant fraction, combined glycerol fraction, the resuspended pellet sample, or the mixed supernatant and pellet sample through a layer of stretched parafilm membrane for 24 h. As shown in Fig 2D and 2E, NSvc2 (red) and RSV virion (green) were detected together in the epithelial cells of SBPHs fed on the mixture of sucrose and the combined glycerol fraction (Fig 2D, row 2). RSV virion was, however, not detected in the microvillus of SBPHs fed on the mixtures of sucrose and the combined supernatant fraction or the resuspended pellet sample (Fig 2D, row 1 and 3). In contrast, when insects were allowed to feed on a mixture of sucrose and the combined supernatant fraction plus the resuspended pellet sample, both NSvc2 and RSV virion were detected in the epithelial cells (OC value = 0.95 ± 0.03; Fig 2D, row 4), similar to that found for the SBPHs fed on the mixture of sucrose and the combined glycerol fraction (OC value = 0.97 ± 0.04). Statistical analysis of RSV transmission using SBPHs fed on various samples also indicated that feeding on the mixture of sucrose and the combined supernatant fraction plus the resuspended pellet sample allowed RSV virion entrance into the midgut epithelial cells for a successful virus transmission (Fig 2E). This finding suggests that RSV NSvc2 is a critical factor mediating RSV virion entrance into SBPH midgut cells. To validate the interaction between RSV NP and NSvc2, we performed yeast two-hybrid assays. Results showed that RSV NP interacted with both NSvc2-N and NSvc2-C (Fig 2F). All these data suggested that NSvc2 protein is critical for RSV virion entrance into SBPH midgut.

### Recombinant amino-terminal soluble region of NSvc2 directly binds to midgut and inhibits subsequent RSV acquisition by SBPH

Previous studies have shown that NSvc2 can be further processed into two mature glycoproteins, namely amino-terminal and carboxyl-terminal NSvc2 [40]. S1A Fig illustrated the predicted structure of NSvc2 and the positions of its signal peptides, transmembrane regions, and the predicted glycan sites. To examine the potential roles of amino-terminal NSvc2 (NSvc2-N) in RSV transmission, we expressed the soluble NSvc2-N protein (referenced to as NSvc2-N:S) in Sf9 insect cells using a recombinant baculovirus expression system (S1B Fig). After purification using the Ni-NTA agarose, the expression of the recombinant NSvc2-N:S was confirmed by Western blot assay using an anti-NSvc2-N polyclonal antibody (Fig 3A).

**Fig 3.**
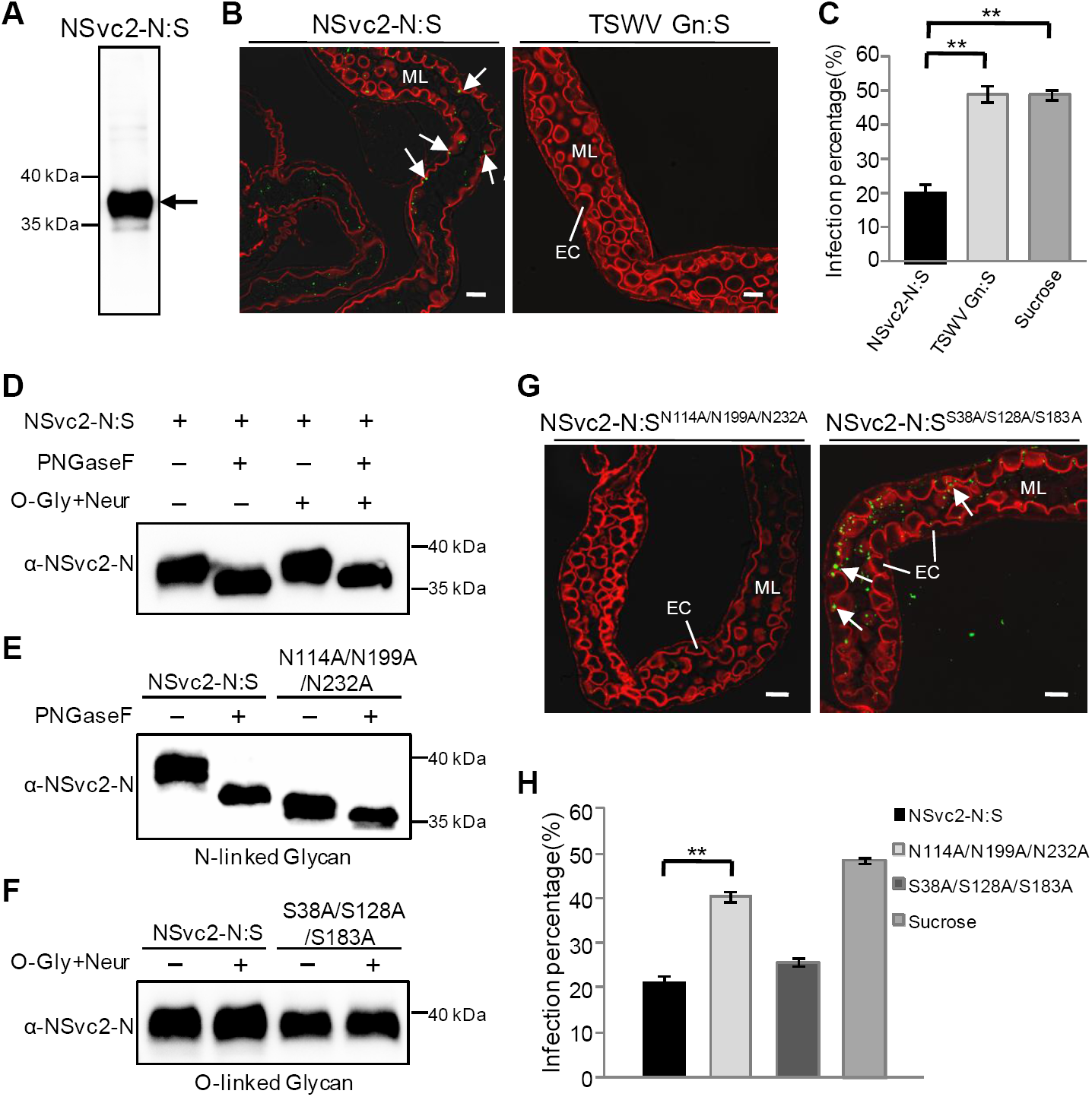
N-glycosylation of NSvc2-N is essential for the recognition of midgut surface receptor. (A) Immunoblot analysis of NSvc2-N:S expression in Sf9 cells. Sizes of protein markers are indicated on the left and the NSvc2-N:S protein band is indicated with an arrow. (B) Binding of purified NSvc2-N:S to microvillus surface of SBPH midgut. Arrows indicate the accumulation of NSvc2-N:S (green, left panel) in midgut lumen. Purified TSWV Gn:S was used as a negative control in this study (right panel). ML, midgut lumen; EC, epithelial cell; Bar, 25 μm. (C) Percentages of RSV acquisition by SBPHs pre-fed with NSvc2-N:S, TSWV Gn:S, or sucrose only. All experiments were performed three times (50 SBPHs for each group). ** *p* < 0.01 by *t*-test analysis. (D–F) Enzymatic de-glycosylation of NSvc2-N:S. Purified NSvc2-N:S was incubated with PNGaseF or O-Glycosidase + Neuraminidase (O-Gly + Neur) to determine the types of glycans (D). PNGaseF was used to remove the N-linked glycans, and O-Gly + Neur were used to remove the O-linked glycans. N-glycosylation (E) and O-glycosylation (F) of the triple asparagine mutant (NSvc2-N:S^N114A/N199A/N232A^) or triple serine mutant (NSvc2-N:S^S38A/S128A/S183A^) were conducted as described for NSvc2-N:S. The NSvc2-N:S^N114A/N199A/N232A^ mutant was treated with PNGaseF while the NSvc2-N:S^S38A/S128A/S183A^ mutant was treated with O-Gly + Neur. (G) The NSvc2-N:S^N114A/N199A/N232A^ mutant failed to bind SBPH midgut (left) but the O-glycosylated NSvc2-N:S^S38A/S128A/S183A^ mutant did (right). ML, midgut lumen; EC, epithelial cell; Bar, 25 μm. (H) RSV acquisition by SBPHs after pre-feeding with different protein samples. The experiment was repeated three times with 50 SBPHs each group. **, *p* < 0.01 by *t*-test analysis.

To further determine whether the recombinant NSvc2-N:S protein can bind midgut epidermal microvillus, SBPHs were allowed to feed on purified NSvc2-N:S for 3 h followed by a 12 h feeding on a sucrose solution to remove unbound NSvc2-N:S. Results of immunofluorescence analyses showed that NSvc2-N:S (green signal) could be readily detected in the midgut lumen near the surface of epithelial cells in the alimentary canal (Fig 3B). As a negative control, SBPHs were allowed to feed on *Tomato spotted wilt virus* (TSWV) encoded glycoprotein (Gn:S), known to bind thrip midgets [28]. As expected, the TSWV Gn:S (green) was not detected in SBPH midguts.

Based on the above results, we further hypothesized that the pre-acquired NSvc2-N:S could prevent RSV acquisition by blocking midgut RSV specific receptors. To test this hypothesis, SBPHs were allowed to feed on the purified NSvc2-N:S for 24 h and then on RSV-infected rice plants for 48 h. The alimentary canals were dissected from SBPHs and probed using the RSV NP or NSvc2-N specific antibodies. Under the confocal microscope, the labeled RSV virion was found in the midgut lumen of SBPHs pre-fed with purified NSvc2-N:S (S2A Fig), suggesting that RSV virion was prevented from entering into the midgut epithelial cells. In contrast, RSV virion was detected in the midgut epithelial cells after the insects were pre-fed with TSWV Gn:S or with sucrose alone (S2B Fig and S2C Fig).

To further confirm the role of NSvc2-N during SBPH acquisition of RSV, SBPHs pre-fed with NSvc2-N:S were allowed to feed on RSV-infected rice plants for 48 h and then on healthy rice seedlings for 12 days. After this feeding period, the insects were tested for RSV infection by ELISA assay. Results showed that pre-feeding SBPHs with NSvc2-N:S did significantly reduce the rate of RSV infection compared with the insects pre-fed with TSWV Gn:S or sucrose only (Fig 3C). This finding indicated that NSvc2-N:S could inhibit RSV entrance into SBPH midgut.

### N-glycosylation of NSvc2-N is required for midgut receptor recognition of RSV

Computer-assisted modeling suggested that NSvc2 might be modified through glycosylation (S1A Fig). To confirm this prediction, purified NSvc2-N:S was incubated with PNGaseF (a N-glycosidase) or O-Glycosidases and Neuraminidase (O-Gly + Neur) to remove the N- or O-linked glycans, respectively. Subsequent SDS-PAGE and immunoblotting analyses showed that the purified NSvc2-N:S protein band was shifted in the gel after the PNGaseF treatment, compared with the non-treated NSvc2-N:S (Fig 3D, compare lane1 and 2). No clear band shift was detected when NSvc2-N:S was treated with O-Gly + Neur (Fig 3D, compare lane1 and 3). When NSvc2-N:S was treated with PNGaseF together with O-Gly + Neur, the protein band shifted as that treated with PNGaseF (Fig 3D, compare lane2 and 4), indicating that NSvc2-N was modified by the N-linked glycans.

We then generated a NSvc2-N:S site-directed alanine-substitution mutant, NSvc2-N:S^N114A/N199A/N232A^ (N114A/N199A/N232A), at its putative N-linked glycan sites, and a mutant, NSvc2-N:S^S38A/S128A/S183A^ (S38A/S128A/S183A), at its putative O-linked glycan sites. These two mutants were purified as describe above for the wild-type (WT) NSvc2-N:S followed by the enzymatic deglycosylation analyses. Results showed that, without PNGaseF treatment, the NSvc2-N:S^N114A/N199A/N232A^ mutant displayed a similar protein band shift in the gel as that shown by the WT NSvc2-N:S treated with PNGaseF (Fig 3E, compare lane 2 and 3). A slight band shift was noticed for the NSvc2-N:S^N114A/N199A/N232A^ mutant, without or with PNGaseF treatment (Fig 3E, compare lane 3 and 4). No O-linked glycan modification was detected for the NSvc2-N:S^S38A/S128A/S183A^ mutant (S38A/S128A/S183A, Fig 3F). This result indicated that residue N114, N199 and N232 of NSvc2-N:S are indeed the N-glycosylation sites.

To investigate whether the N-linked glycosylation can affect the recognition of NSvc2-N:S by midgut surface receptors, SBPHs were fed with the two mutant proteins, respectively, for 3 h followed by a 12 h feeding on a sucrose only solution to clean insect alimentary canals. Insects fed with the NSvc2-N:S^N114A/N199A/N232A^ mutant protein showed almost no green labeling signal at the surface of midgut microvillus. In contrast, insects fed with the NSvc2-N:S^S38A/S128A/S183A^ mutant protein did (Fig 3G). ELISA results showed that the NSvc2-N:S^N114A/N199A/N232A^ mutant protein had no obvious effect on SBPH RSV acquisition (Fig 3H). Consequently, we conclude that modification of NSvc2-N protein through N-glycosylation is important for midgut surface receptor recognition.

### RSV virion:NSvc2-N:NSvc2-C complexes enter into the endosomes and NSvc2-C separates from RSV virion:NSvc2-N after releasing from endosomes

Endocytosis is an important process for several circulative-transmitted animal viruses during entering into animal cells [43]. The localization of NSvc2-N and RSV virion suggested that RSV virion and NSvc2-N has entered into endosomal-like vesicle (Fig 1*B* and 1F). To further investigate the localization of RSV virion and NSvc2-N after recognition by SBPH midgut cell receptor(s), early and late endosome specific markers (Rab5, EEA1 and Rab7) were used to visualize these vesicles [44, 45]. Results showed that RSV virion did co-localize with the early endosome Rab5 marker and the late endosome Rab7 marker (Fig 4A and 4B). In the same study, NSvc2-N and carboxyl-terminal protein of NSvc2 (NSvc2-C) were also found to co-localize with early endosome EEA1 marker (Fig 4C and 4D).

**Fig 4.**
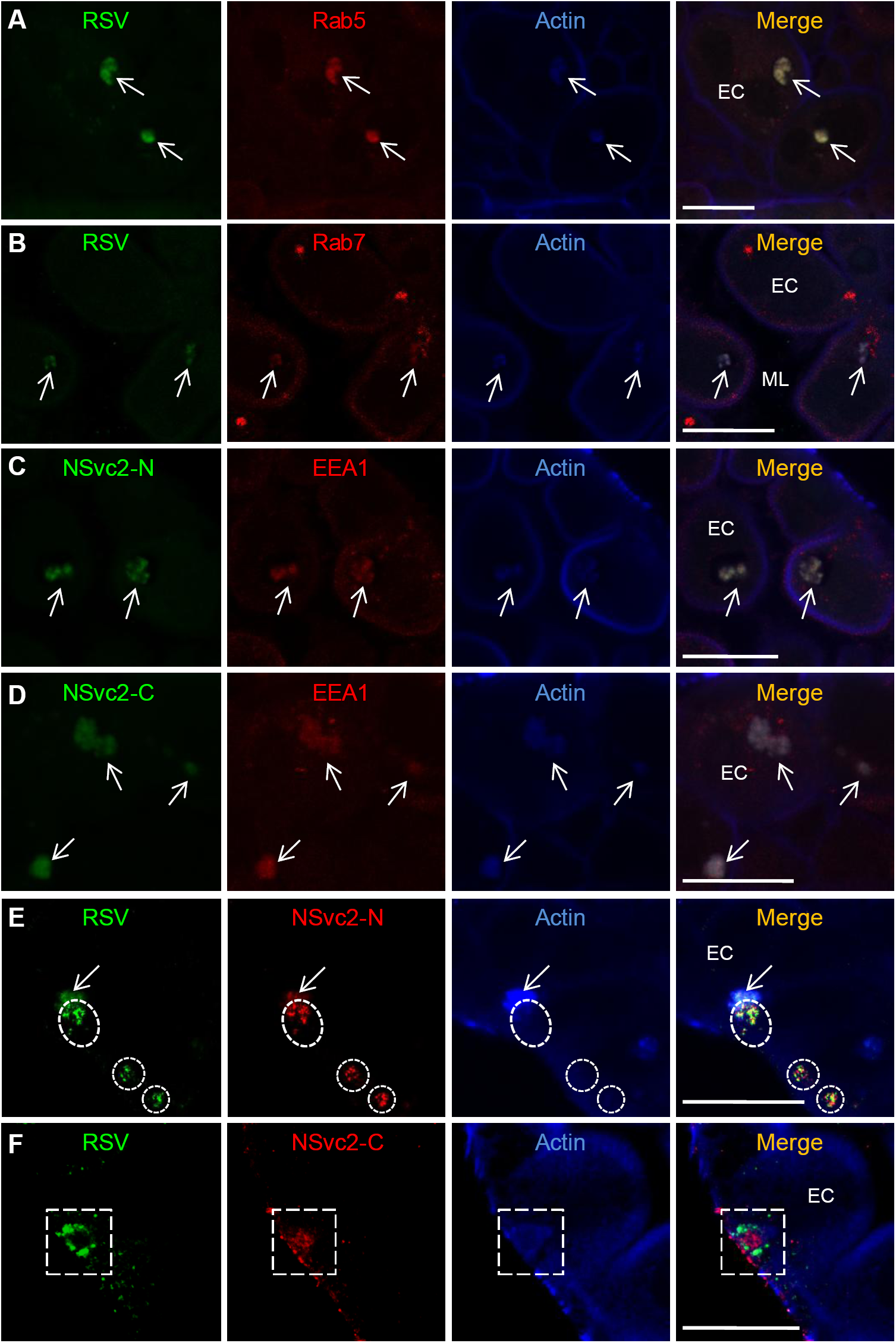
RSV virion, NSvc2-N and NSvc2-C are co-localized with early or late endosomes, and NSvc2-C separates from RSV virion and NSvc2-N complexes after releasing from endosomes in midgut epithelial cells. (A and B) RSV virion was detected in early endosomes labeled with the Rab5 antibody (A) or in late endosomes labeled with the Rab7 antibody (B) in midgut epithelial cells. Co-localizations of RSV virion with Rab5 or Rab7 or actin in different endosomes are indicated by arrows. ML, midgut lumen; EC, epithelial cell; Bar, 25 μm. (C and D) NSvc2-N and NSvc2-C were detected in early endosomes labeled with the EEA1 antibody in epithelial cells. Co-localizations of NSvc2-N or NSvc2-C with EEA1 or actin in different endosomes are indicated by arrows. Bar, 25 μm. (E) RSV virion and NSvc2-N were released from endosome into the cytosol of epithelial cells. The white dashed cycles indicated a region that RSV and NSvc2-N were detected in the cytosol while actin-labeled endosome was absent. Bar, 25 μm. (F) RSV virion but not NSvc2-C was released from the late endosome in epithelial cells. The white dashed boxes indicated a region that RSV virion was just released from the endosome while NSvc2-C proteins were still associated with actin-labeled endosome. Bar, 25μm.

To examine the potential role(s) of NSvc2-C, SBPHs were allowed to feed on RSV-infected rice plants for 4, 8, 16 or 24 h, and then used for immunofluorescence labeling assays. NSvc2-C with red labeling signal was observed together with RSV virion (green) at the surface of microvillus (blue) (S3A Fig and S3E Fig; OC value = 0.86 ± 0.04), and in the endosomal-like vesicles in epithelial cells (S3B Fig and S3F Fig; OC value = 0.95 ± 0.03) at 4 and 8 h post feeding. At 16 and 24 h post feeding, RSV virion (green) had accumulated alone in the cytoplasm of epithelial cells (S3C Fig and S3G Fig [OC value = 0.18 ± 0.02], and S3D Fig and S3H Fig [OC value = 0.32 ± 0.06]), suggesting that NSvc2-C remained inside the endosomal-like vesicles while RSV virion was released from the endosomal-like vesicles and accumulated in the cytoplasm. To confirm this finding, the localization of RSV virion, NSvc2-N and NSvc2-C was examined just at the time that RSV virion release from endosomes. The results showed that after being released from endosomes, NSvc2-N continued to associate with RSV virion in cytosol (Fig 4E, white dashed circles), whereas NSvc2-C stayed inside the actin-labeled endosome and was not released into cytosol of epithelial cells (Fig 4F, white dashed box). This finding indicates that NSvc2-C associates with RSV virion:NSvc2-N in endosome but was separated from them after the virion were released from endosome.

### NSvc2-C induces insect cell membrane fusion under acidic conditions

To determine whether NSvc2-N and/or NSvc2-C play roles in membrane fusion, we fused a signal peptide of baculovirus (gp64) to NSvc2-N and NSvc2-C, and expressed these proteins individually in insect *Spodoptera frugiperda* (Sf9) cells by the recombinant baculovirus expression system (Fig 5A). Expressions of these recombinant proteins were confirmed by an immunolabeling assay. Under the laser scanning confocal microscope, red labeling fluorescence signal representing NSvc2-N or NSvc2-C was observed on the Sf9 cell membranes (Fig 5B). We then tested whether NSvc2-N or NSvc2-C could trigger cell membrane fusion under acidic conditions. Sf9 cells infected with the recombinant NSvc2-N or NSvc2-C baculovirus were treated with a PBS, pH 5.0, for 2 min and then grown in a pH neutral medium. Numerous cell-cell fusions (syncytium) were observed at 4 h post acidic PBS treatment of Sf9 cells infected with the NSvc2-C baculovirus (Fig 5E and 5G). In contrast, no significant cell-cell fusion was observed for cells infected with the NSvc2-N or empty baculovirus (Fig 5C and 5D). In addition, cell membrane fusion was observed in cells co-infected with NSvc2-N and NSvc2-C baculoviruses (Fig 5F and 5G), confirming that RSV NSvc2-C, but not NSvc2-N, played an important role in cell-cell fusion under acidic conditions.

**Fig 5.**
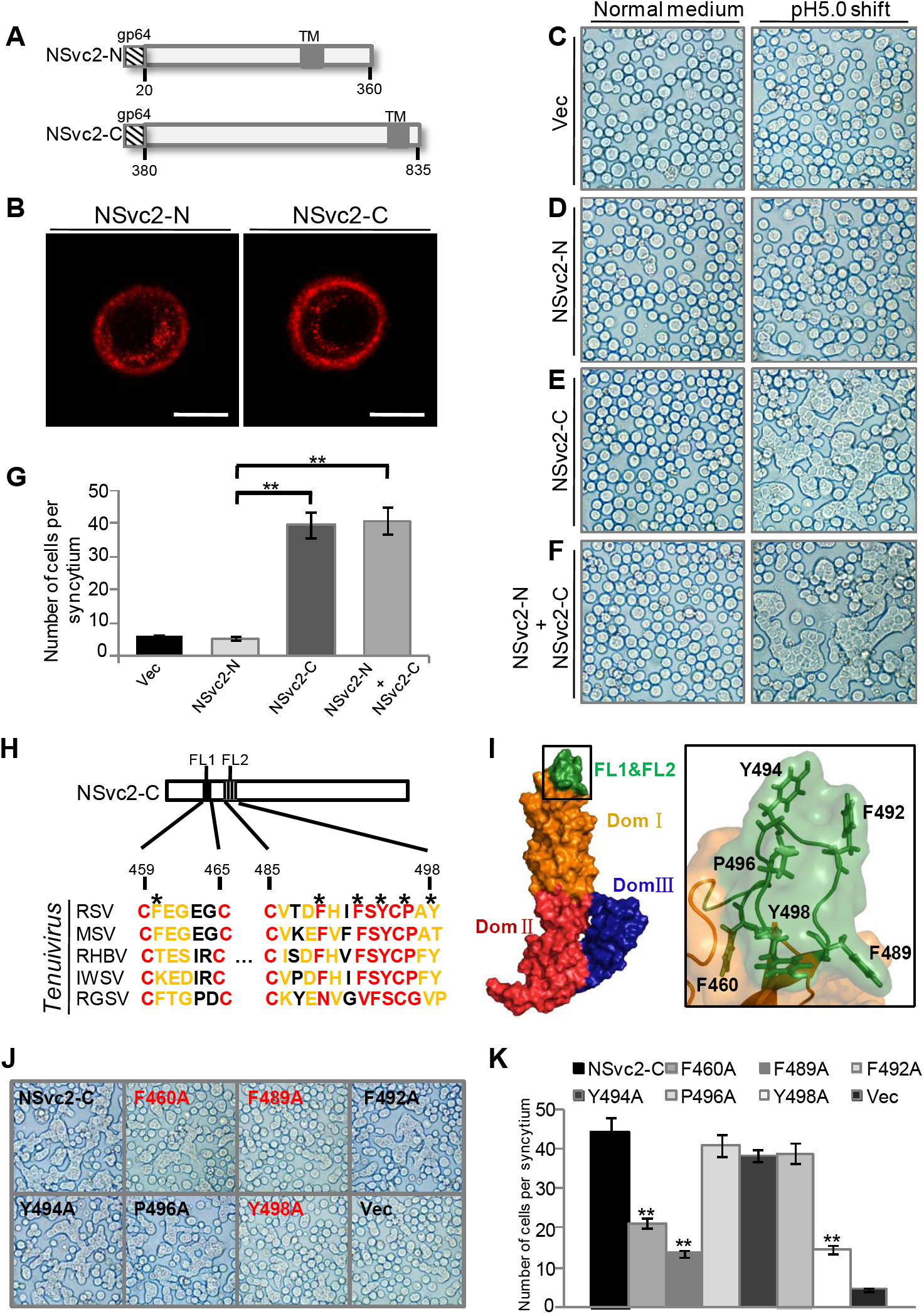
NSvc2-C hydrophobic fusion-loop motifs are required for cell-cell membrane fusion. (A) Schematic diagrams showing recombinant NSvc2-N and NSvc2-C. Baculovirus gp64 signal peptide was used to replace the original signal peptide at the N-terminus of NSvc2-N or NSvc2-C. (B) Immunofluorescence labeling of the recombinant NSvc2-N or NSvc2-C in Sf9 cells. Sf9 cells were infected with baculoviruses carrying the recombinant NSvc2-N or NSvc2-C (MOI=5). The infected cells were probed with the NSvc2-N or NSvc2-C specific antibody at 48 h post infection followed by a TRITC fluorescent-labeled secondary antibody. Bar, 10 μm. (C–F) Fusogenic activity assays using Sf9 cells expressing the recombinant NSvc2-N, NSvc2-C or both (NSvc2-N + NSvc2-C). The Sf9 cells were infected with recombinant baculoviruses (MOI=5). After 48 h post infection, and the growth medium was replaced with a PBS, pH 5.0, for 2 min and then changed back to the normal growth medium for 4 h. Formation of syncytium was observed under a microscope. (G) The number of cells showing syncytium was counted under the microscope. The experiment was repeated three times. **, *p* < 0.01 by the student *t*-test analysis. (H) Sequence alignment using the two fusion loop sequences (FL1 and FL2) from NSvc2-C of five different tenuiviruses (RSV, MSV, RHBV, IWSV and RGSV). Red residues are highly conserved and yellow residues are semi-conserved. Residues indicated with asterisks are hydrophobic residues and were selected for the site-directed mutagenesis. The resulting mutants were later tested for their abilities to induce cell-cell membrane fusion. (I) Two fusion loops (FL1 and FL2, green) were found at the top of the 3D-structure model of NSvc2-C. Three different domains are shown in three different colors, and the boxed region on the 3D-structure is enlarged and shown on the right side. Locations of the hydrophobic residues are displayed. (J and K) Analyses of the fusogenic activities caused by the WT or mutant NSvc2-C. Sf9 cells were infected with recombinant baculoviruses carrying the WT or mutant NSvc2-C for membrane fusion assay (J). The number of cells showing syncytium was recorded and analyzed (K). The experiments were repeated three times. **, *p* < 0.01.

### Conserved hydrophobic fusion-loop motifs in NSvc2-C are crucial for fusogenic activity

To further investigate the function of NSvc2-C in cell membrane fusion, we generated a three-dimensional (3D) structure of NSvc2-C through a homology modeling approach (S4A Fig and S4B Fig). This 3D structure consisted of three distinct domains: domain I (yellow), II (red), and III (blue). Two putative fusion loops (green) were found at the top of the 3D structure. Similar fusion loops were reported to be responsible for cell-cell fusion during animal virus infections [46, 47]. In this study, we constructed three NSvc2-C fusion domain deletion mutants (i.e., ΔFL1, ΔFL2 and ΔFL1+ΔFL2), and expressed them individually in Sf9 cells followed by immunoblotting (S4C Fig and S4D Fig). The fusogenic activities of these deletion mutants were then examined in Sf9 cells using the recombinant baculovirus expression system. Results showed that the number of syncytial cells induced by the ΔFL1 or ΔFL2 mutant was much less than that induced by the WT NSvc2-C (S4E Fig and S4F Fig). The lowest number of syncytial cells was, however, observed in the Sf9 cells infected with the baculovirus carrying the ΔFL1+ΔFL2 double deletion mutant. Based on this finding, we conclude that the two NSvc2-C fusion loops play critical roles in cell-cell fusion.

To identify the amino acid residue(s) important for fusogenic activity, we aligned the RSV fusion loop sequences (Cys459-Cys465, Loop1 and Cys485-Tyr498, Loop 2) with the loop sequences of other four tenuiviruses (Fig 5H). The alignment result indicated that these two fusion loops were relatively conserved among the five tenuiviruses. Six conserved hydrophobic amino acid residues (Phe460, Phe489, Phe492, Tyr494, Pro496 and Tyr498) were found at the unique vertex in the modeled NSvc2-C 3D structure (Fig 5I). Introduction of mutations into these six amino acid residues suggested that three residues (F460A, F489A, and Y498A) were important for membrane fusion activity (Fig 5J and 5K).

### NSvc2-C fusogenic-activity-deficient mutant fails to mediate RSV virion release from endosome

The above results showed that RSV NSvc2-N and NSvc2-C had different functions during RSV transmission via SBPH. To elucidate the role of NSvc2-C in RSV entrance into midgut cells during virus acquisition, we analyzed the fusion loop mutants for their functions in RSV virion release from endosome. Purified RSV virion was incubated for 3 h with Sf9 cell crude extracts containing the full length NSvc2 and then used the mixture to feed SBPHs for 24 h. Midguts were isolated from SBPHs and probed for the presence of RSV virion and NSvc2-N through immunofluorescence labeling. Result showed that RSV virion (green) and NSvc2-N (red) were both present in the endosomal-like vesicles in epithelial cells, indicating that both RSV virion and NSvc2-N had entered into the midgut cells (Fig 6A, upper row). Result also showed that RSV virion was released from endosome at 24 h post feeding. We then prepared Sf9 cell crude extracts containing the N-glycosylation-deficient mutant NSvc2^N114A/N199A/N232A^ or the fusogenic-activity-deficient mutant NSvc2^F460A/F489A/Y498A^. Using the same feeding method described above, RSV virion was detected in the midgut lumen but not in the epithelial cells of SBPHs fed with the mixture of RSV virion and NSvc2^N114A/N199A/N232A^ mutant (Fig 6A, middle row). This study also showed that RSV virion and NSvc2^F460A/F489A/Y498A^ mutant overcame the midgut barrier and entered together into the endosomal-like structures. However, RSV virion and NSvc2^F460A/F489A/Y498A^ mutant were hardly detected in the cytoplasm of epithelial cells. Also, greater number of endosomal-like structures was found in epithelial cells of SBPHs fed with the mixture of RSV virion and NSvc2^F460A/F489A/Y498A^ mutant than that fed with the mixture of RSV virion and wild type NSvc2 (Fig 6A, bottom row). Statistical analysis further confirmed that RSV virion did enter endosomes but failed to be released into cytosol after SBPHs were fed with the mixtures of purified RSV virion and NSvc2^F460A/F489A/Y498A^ mutant, while RSV virion did not enter endosomes when SBPHs were fed with the mixtures of purified RSV virion and NSvc2^N114A/N199A/N232A^ mutant (Fig 6B). This finding indicates that the fusogenic-activity-deficient NSvc2 mutant can mediate virion passage through epithelial cells but failed to release RSV virion from endosomal-like vesicles.

**Fig 6.**
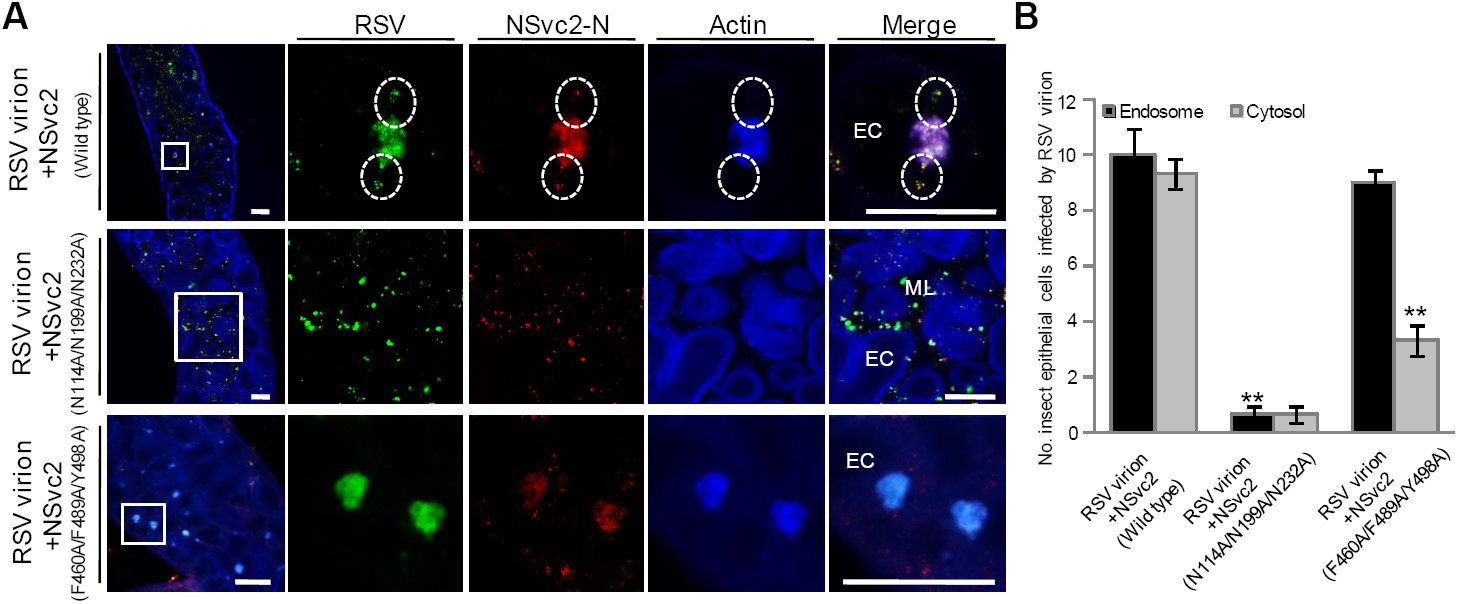
NSvc2 functions as a molecular bridge to mediate RSV virion entrance into SBPH midgut cell. (A) Crude extract of Sf9 cell expressing WT NSvc2, NSvc2^N114A/N199A/N232A^ or NSvc2^F460A/F489A/Y498A^ was incubated with purified RSV virion for 3 h at 4 °C, and then used to feed SBPHs for 24 h. Insect midguts were dissected and detected for RSV virion and NSvc2-N using specific antibodies. The boxed regions in the left column were enlarged and shown in the second to fifth columns on the right. The white circled areas show the release of RSV virion (green) and NSvc2-N (red) from endosome into cytosol. The actin labeling signal is shown in blue. ML, midgut lumen; EC, epithelial cell; Bar, 25 μm. (B) Statistical analysis of RSV infection in SBPH epithelial cells after feeding on the mixtures containing purified RSV virion and the WT or mutant NSvc2-C. Each bar represents three independent biological repeats from each experiment with 50 SBPHs per treatment. **, *p* < 0.01 by student *t*-test analysis.

**Fig 7.**
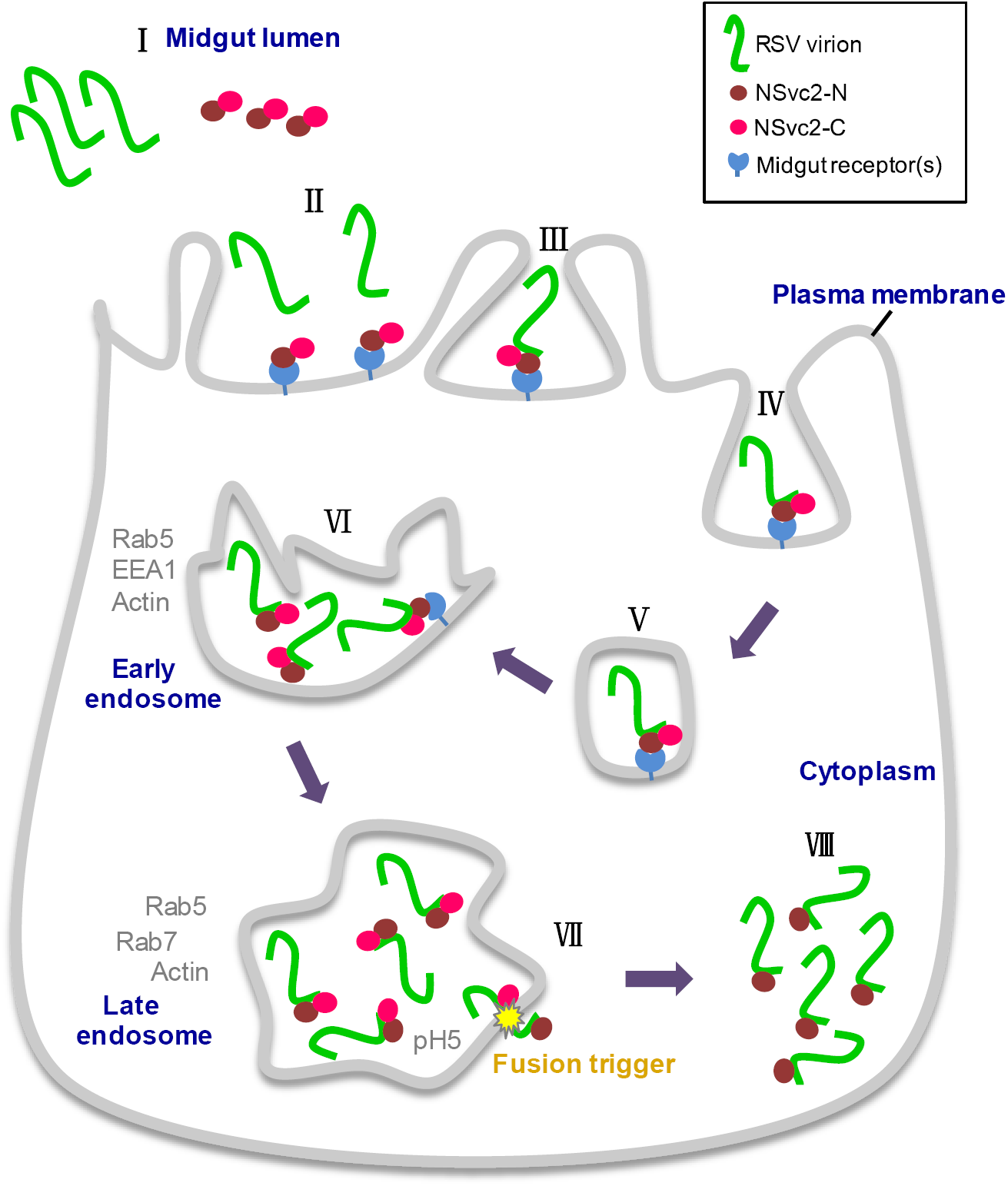
Working model for RSV virion entrance into SBPH midgut cells. During SBPH feeding on RSV-infected rice plants, RSV virion, NSvc2-N and NSvc2-C are acquired by the vector and enter insect midgut lumen (Step I). After reaching microvillus surface of midgut lumen, NSvc2-N protein directs the virion:NSvc2-N:NSvc2-C complexes to microvillus surface through recognizing the unidentified microvillus surface receptor(s) (Steps II and III). The complexes-attached midgut lumen membrane undergoes endocytosis and compartmentalizes RSV virion, NSvc2-N and NSvc2-C in vesicles (Steps IV and V). These vesicles further develop into early endosomes (Step VI) and then late endosomes (Step VII). The acidic condition inside the late endosomes triggers NSvc2-C to induce cell-cell membrane fusion (Step VII). Finally, RSV virion:NSvc2-N complexes are released from the late endosomes into the cytosol of epithelial cells for further RSV spread in midgut cells (Step VIII).

## Discussion

Understanding how plant viruses are transmitted through their insect vectors in field is one of the key steps to manage virus diseases worldwide. Insect midgut is a major barrier to block the entrance of non-compatible plant viruses. In this study, we used RSV and its SBPH vector as a working model to elucidate the molecular mechanism controlling RSV virion entrance into SBPH midgut for replication. We demonstrated here for the first time that RSV, a circulative and propagative transmitted *Tenuivirus*, used a molecular bridge strategy to overcome the midgut barrier in SBPH for virus persistent transmission.

In this study, we confirmed that glycoprotein NSvc2 of RSV is not a structural protein of RSV virion, and found that in the absence of this glycoprotein, RSV virion is unable to overcome the midgut barrier (Fig 2). RSV glycoprotein NSvc2 was further processed into an amino-terminal part protein known as NSvc2-N and a carboxyl-terminal part protein known as NSvc2-C in RSV-infected cells [39, 40]. We found that NSvc2-N accumulated on midgut surface during virus acquisition by SBPH. We also found that the ectopically expressed and purified soluble NSvc2-N:S bound directly to midgut surface and inhibited the subsequent RSV acquisition by SBPH. We consider that this soluble NSvc2-N:S protein can recognize SBPH midgut surface receptor(s). Our enzymatic deglycosylation result showed that NSvc2-N could be modified by N-linked but not O-linked glycans. The glycan-modification of NSvc2-N might be different from the N- and O-glycosylation of TSWV Gn protein reported previously [28]. It is noteworthy that the N-glycosylation-deficient NSvc2-N^N114A/N199A/N232A^ mutant was unable to interact with midgut surface and to block RSV acquisition by SBPH. Moreover, feeding SBPHs with a mixture of purified RSV virion and crude extract from Sf9 cells expressing full length NSvc2 resulted in a successful transmission of RSV. In contrast, the N-glycosylation-deficient NSvc2 mutant was unable to facilitate RSV virion to pass through SBPH midgut barrier. Therefore, we propose that N-glycosylation modification plays an important role in interaction between RSV virion and SBPH midgut receptor(s). A sugar transporter in SBPH midgut cell was recently found to play critical role in RSV transmission [48]. It would be interested in investigating the interactions between NSvc2 and sugar transporter protein and the biological function of glycosylation for these interactions in the future.

Although the function of RSV glycoprotein NSvc2 was proposed to be similar to that of tospovirus glycoprotein or phytoreovirus spike protein during virus acquisition and transmission [29, 49], NSvc2 is not present on the surface of non-enveloped filamentous RSV virion while both tospovirus glycoprotein and phytoreovirus spike protein are located virion surface. Purified RSV virion is not capable of entering SBPH midgut cells. This inability can, however, be rescued by addition of glycoprotein NSvc2 (Fig 2D and Fig 6A). For non-persistent or semi-persistent transmitted plant viruses, the helper component proteins or helper factors were shown to act as molecular bridges for the interactions between virion and the cuticle of their insect vectors [4, 6]. Although these helper component proteins or factors are not located on the surface of purified virion, these factors are absolutely required for insect transmissions [18, 50]. Our results presented in this paper indicate that this helper factor theory can also be used to explain the function of glycoprotein NSvc2 during RSV circulative and propagative transmission by SBPH. First, the RSV glycoprotein NSvc2 is not a virion structural protein (Fig 2). Second, the glycoprotein NSvc2 acts as a molecular bridge to ensure the interaction between RSV virion and SBPH midgut receptor(s) (Fig 3). Third, the glycoprotein NSvc2 is absolutely required for RSV virion entrance into SBPH midgut cells (Fig 2 and Fig 6). These findings strongly supported that RSV glycoprotein NSvc2 is a helper factor protein and this helper factor functions as a molecular bridge to allow the proper interaction between RSV viroin and SBPH midgut during persistent transmission of *Tenuiviru*s.

After recognition and interaction with RSV virion, midgut epithelial cells underwent endocytosis, resulting in an enclosure of RSV virion:NSvc2 complexes in early and then late endosomes. After that, RSV virion was released from the late endosomes into cytoplasm for replication and for further spreading to other cells. NSvc2-C was initially found together with RSV virion on midgut surface, and later in endosomal-like vesicles. Although NSvc2-C retained inside the endosomes, NSvc2-N-associated RSV virion was released into cytoplasm. We determined that, under acidic conditions, the expressed NSvc2-C could cause Sf9 cell membrane to fuse. To identify the amino acid residue(s) controlling this membrane fusion activity, a homology based modeling approach was used to create a 3D structure of NSvc2-C followed by site-directed mutagenesis. With the aid of homology modeling, we identified that NSvc2-C residue F460, F489 and Y498 played critical roles during cell membrane fusion. Importantly, the NSvc2 mutant that failed to cause cell membrane fusion was unable to release RSV virion from endosomes into cytoplasm. It is likely that the acidic condition inside endosomes caused conformation change of NSvc2-C, leading to cell membrane fusion. The two fusion loops found in NSvc2 are highly conserved among the members in the genus *Tenuivirus* and are critical for cell membrane fusion induction.

Current gene function studies for plant multi-segmented negative-strand RNA viruses are difficult due mainly to the lack of proper reverse genetics methods. In this paper, we described a new and useful approach to overcome this obstacle. We first expressed the WT or mutant NSvc2 individually in Sf9 insect cells and then incubated the Sf9 cell crude extracts with purified RSV virion. After feeding SBPHs with a WT or mutant NSvc2 extract mixed with purified RSV virion, we successfully determined that virion from the sample containing the WT NSvc2 and purified RSV virion overcame the midgut barrier, entered the epithelial cells, and then released from late endosomes to cytosol for further replication and transmission. The defective NSvc2 mutants failed to interact with midgut surface and thus were unable to mediate the entrance of RSV virion into epithelial cells. In addition, using this approach, we were able to confirm that the cell membrane fusion defective mutant could not mediate the release of RSV virion from endosomes to cytosol. The assay methods developed in this study should benefit gene function studies for viruses whose infectious clones are currently difficult to make. We proposed that this newly developed technology can not only be used to investigate the functions of tenuivirus glycoproteins, but can also be used to determine the functions of glycoproteins encoded by other plant multi-segmented negative-strand RNA viruses.

Taken together, we conclude that circulative and propagative transmitted RSV uses a molecular bridge strategy to bring RSV virion to SBPH midgut surface during vector transmission. Based on the findings presented in this study, we propose a working model for plant tenuiviruses (Fig 6C). In this model, the NSvc2-N and NSvc2-C do not associate with RSV virion in infected plant cells. After plant sap containing RSV virion, NSvc2-N, and NSvc2-C is acquired into the midgut lumen of SBPH, NSvc2-N recognizes the unidentified midgut cell surface receptor(s) and acts as a molecular bridge to ensure the interaction between RSV virion and midgut surface. Upon attachment of RSV virion:NSvc2-N:NSvc2-C complexes to midgut surface receptor(s), midgut cells undergo endocytosis, resulting in compartmentalization of RSV virion, NSvc2-N, and NSvc2-C in early and then late endosomes. The acidic condition inside the late endosomes triggers a conformation change of NSvc2-C, and the conformation-changed NSvc2-C cause cell membrane to fuse. Finally, the RSV virion:NSvc2-N complexes are released into cytosol (Fig 6C). Findings presented in this paper demonstrate a new type of virus–vector midgut interaction that requires a virally encoded molecular bridge during virus persistent transmission. This type of mechanism has never been shown for persistent transmitted plant viruses or animal viruses. This new finding expands our understanding of molecular mechanism(s) controlling virus–insect vector interactions during virus transmission in nature.

## Materials and methods

### Insect and virus maintenance

*Rice stripe tenuivirus* (RSV) was previously isolated from an RSV-infected rice plant, and maintained inside a growth chamber at the Jiangsu Academy of Agricultural Sciences, Jiangsu Province, China. Small brown planthopper (SBPH) was reared on seedlings of rice cv. Wuyujing NO.3 inside growth incubators set at 26.5 °C and a photoperiod of 16 h / 8 h (light / dark). Rice seedlings were changed once every 12 days to ensure sufficient nutrition as described [51].s

### Isolation of RSV virion

Eight hundred milliliter precooled 0.1 M phosphate buffer (PB), pH7.5, with 0.01 M EDTA was added to 50 g RSV-infected rice leaf tissues followed by 5 min homogenization in a blender. The homogenate was centrifuged at 8,000 × *g* for 20 min at 4°C. The resulting supernatant was slowly mixed with 6 % PEG 6000 and 0.1 M NaCL, and stirred overnight at 4 °C. After centrifugation at 8,000 × *g* for 20 min, the pellet was resuspended in 0.01 M PB and then centrifuged again at 150,000 × *g* for 2 h. The pellet was resuspended in 6 ml 0.01 M PB buffer and laid on a 4 ml 20 % glycerol cushion inside a centrifuge tube followed by a centrifugation at 150,000 × *g* for 2 h. Different fractions inside the centrifuge tube were collected individually and the pellet was resuspended in PB buffer (contains 30 % glycerol) prior to storage at −70 °C.

### Immunofluorescence staining of SBPH midgut organs

Immunofluorescence staining assay was performed as described previously with specific modifications [52]. Midguts were obtained from second-instar nymphs and fixed overnight in a 4 % paraformaldehyde (PFA, Thermo Fisher, USA) solution at 4 °C. After three rinses in a 0.01 M phosphate-buffered saline (PBS), pH 7.4, the midguts were treated for 30 min in a 2 % Triton X-100 solution followed by 1 h incubation in a specific primary antibody. The midguts were then incubated in a specific fluorescence conjugated secondary antibody or a fluorescence conjugated actin antibody for 2 h at room temperature (RT). The midguts were rinsed three times in PBS and mounted in an antifade solution (Solarbio). The mounted midguts were examined under an inverted Leica TCS SP8 fluorescent confocal microscope (Leica Microsystems, Solms, Germany). LAS X software was used to analyze fluorescence spectra to determine co-localization of two proteins.

Rabbit polyclonal antibody against RSV NSvc2-N or NSvc2-C was produced in our laboratory. Mouse monoclonal antibody against RSV NP was a gift from Professor Jianxiang Wu, Zhejiang University, Hangzhou, China. Primary antibodies also used in this study included Rab5 anti-rabbit IgG (C8B1; Cell Signaling Technology), Rab7 anti-rabbit IgG (D95F2; Cell Signaling Technology), and EEA1 anti-mouse IgG (NBP2-36568; Novus). Secondary antibodies used in this study were FITC conjugated rabbit anti-mouse IgG (F9137; Sigma) or goat anti-rabbit IgG (F9887; Sigma), TRITC conjugated goat anti-rabbit IgG (T6778; Sigma), Alexa Fluor 647 phalloidin (A22287; Invitrogen) or Rhodamine Phalloidin (R415; Invitrogen).

### Recombinant baculovirus expression in insect cell

Construction of recombinant baculovirus was the same as described previously [53]. Sequence encoding RSV NSvc2-N:S (amino acid positions 20 to 265, lacking the signal peptide and the transmembrane domain) was PCR-amplified using a cDNA from a RSV-infected rice plant. The resulting PCR fragment represented the RSV NSvc2-N:S sequence and a six-His tag sequence at its 3’ end terminus. The Gp64 signal peptide sequence was then fused to the 5’ end terminus of NSvc2-N:S sequence via overlapping PCR. The final PCR product was cloned into vector pFastBac1 (S1 Table). The site-directed mutagenized mutants were constructed as described [54]. All plasmids were sequenced, transformed individually into DH10Bac™ cells to generate recombinant baculoviruses as instructed by the manufacturer (Invitrogen). The recombinant baculoviruses were co-transfected individually with the FnGENE HD Transfection Reagent (Promega) into *Spodoptera frugiperda* (Sf9) cells to obtain the stable expression of recombinant baculoviruses.

### Expression and purification of soluble proteins

Sf9 cells were infected with recombinant baculoviruses at a multiplicity of infection (MOI) of five, and the infected-cells were collected at 72 h post infection. The infected-cells were lysed in 10 ml PBS using a ultrasonic cell crusher followed by centrifugation at 4 °C to remove cell debris. After centrifugation, the supernatant was collected, incubated with nickel-nitrilotriacetic acid resin (Ni-NTA, Germany) for 2 h, and then loaded onto a chromatographic column (Bio-Rad Hercules, California). After separation, the column was washed with two bed volumes of 50 mM imidazole in PBS, and the recombinant protein was eluted with 250 mM imidazole followed by dialysis against PBS prior to storage at −70 °C until use.

### Glycosylation and Immunoblotting assays

Purified NSvc2-N:S and various mutant proteins were deglycosylated with PNGaseF as instructed (NEB, USA) to remove N-linked glycans or Neuraminidase and O-Glycosidase (NEB, USA) to remove O-linked glycans. Proteins were mixed with 0.5 % SDS and 40 mM DTT, and incubated at 100°C for 10 min. After denaturation, buffer and glycosidases were added to the samples and incubated for 3 h at 37 °C. The enzyme-treated samples were mixed with a loading buffer containing SDS and boiled for 10 min prior to electrophoresis in 10 % (w/v) SDS-PAGE gels. The separated proteins were transferred to nitrocellulose membranes and the membranes were probed with a polyclonal rabbit antibody against NSvc2-N (1:5,000 dilution) and then a goat anti-rabbit IgG HRP conjugate (31466; Invitrogen). The detection signal was visualized using the ChemiDoc™ Touch Imaging System (Bio-Rad, Hercules, California).

### Glycoprotein midgut binding assays

Midgut binding assays were performed as described previously [28]. Briefly, second-instar nymphs were placed in open-ended EP centrifuge tubes and fed with mixtures of purified glycoproteins resuspended in a TF buffer (PBS with 10 % glycerol, 0.01 % Chicago sky blue, and 5 mg / ml BSA) through a stretched parafilm membrane. After 3 h feeding, the glycoprotein mixtures were replaced with a 10 % sucrose solution for another 12 h feeding to clear midguts, indicated by the disappearance of the Chicago sky blue dye from the midguts. The insects were then dissected, fixed in PFA and analyzed by immunofluorescence staining as described above.

### Detection of SBPH RSV acquisition using Enzyme linked immunosorbent assay (ELISA)

Second-instar nymphs were placed in an empty bottle for 2 h and then allowed to acquire purified protein through a stretched parafilm membrane for 24 h. The pre-fed SBPHs were then allowed to feed on RSV-infected rice seedlings for 48 h prior to feeding on healthy rice seedlings for another 12 days. SBPHs were transferred individually into a centrifuge tube and grinded thoroughly in PBS buffer. After brief centrifugation, individual supernatant samples were blotted on nitrocellulose membranes and the membranes were probed with a 1:5,000 (v/v) diluted monoclonal antibody against RSV NP, and then a 1:20,000 diluted secondary alkaline phosphatase (AP)-coupled goat anti-mouse IgG (Sigma). Detection signal was visualized using a 5-bromo-4-chloro-3-indo-lylphosphate-nitroblue tetrazolium (BCIP-NBT) solution (Sangon Biotech, Shanghai, China). Three independent experiments with 50 nymphs / treatment were performed.

### Yeast two-hybrid assay

Yeast two-hybrid assays were performed according to the instructions from the manufacturer (Clontech, TaKaRa, Japan). Briefly, one bait vector and a specific prey vector, described in the result section, were co-transformed into the Y2HGold strain cells by heat shock method. In addition, vector pGADT7-T was co-transformed into the Y2HGold strain cells with vector pGBKT7-53 or vector pGBKT7-Lam, and used as a positive and a negative control, respectively. All co-transformed cells were first grown on a synthetic dextrose medium lacking Tryptophan and Leucine amino acid (SD-Trp-Leu) for 3 days at 30 °C, and then on a synthetic dextrose medium lacking Tryptophan, Leucine, Histidine and Ademethionine amino acid (SD-Trp-Leu-His-Ade) for 5 days at 30 °C.

### Low pH-induced membrane fusion assay

Sf9 cells were infected with different recombinant baculoviruses at a MOI of five for 48 h. The infected cells were washed twice with fresh medium, treated with a PBS buffer, pH 5.0, for 2 min and then in a pH neutral medium. The cells were incubated for 4 h at 28 °C and then examined for the cell-cell membrane fusion under an Olympus IX71 inverted fluorescence microscope (Olympus, Hamburg, Germany).

### Homology modeling of NSvc2-C

Homology based modeling of NSvc2-C was as described previously [55, 56]. Briefly, RSV NSvc2-C sequence was used to search the I-TASSER Server software. Based on the high TM-score value (0.805), glycoprotein of *Sever Fever with Thrombocytopenia Syndrome Phlebovirus* (SFTSV) (PDB: 5G47) was chosen as the template to build the homology based model of RSV NSvc2-C. Amino acid residues and their surface displays in the three-dimensional structure were predicted using the PyMOL program.

## Acknowledgments

We thank Prof. Yidong Wu (College of Plant Protection, Nanjing Agricultural University, China) for kindly providing the insect Sf9 cell lines. We also thank Prof. Tong Zhou (Institute of Plant Protection, Jiangsu Academy of Agricultural Sciences) for kindly providing the RSV-infected rice samples.

## Supporting information

**Fig S1.**
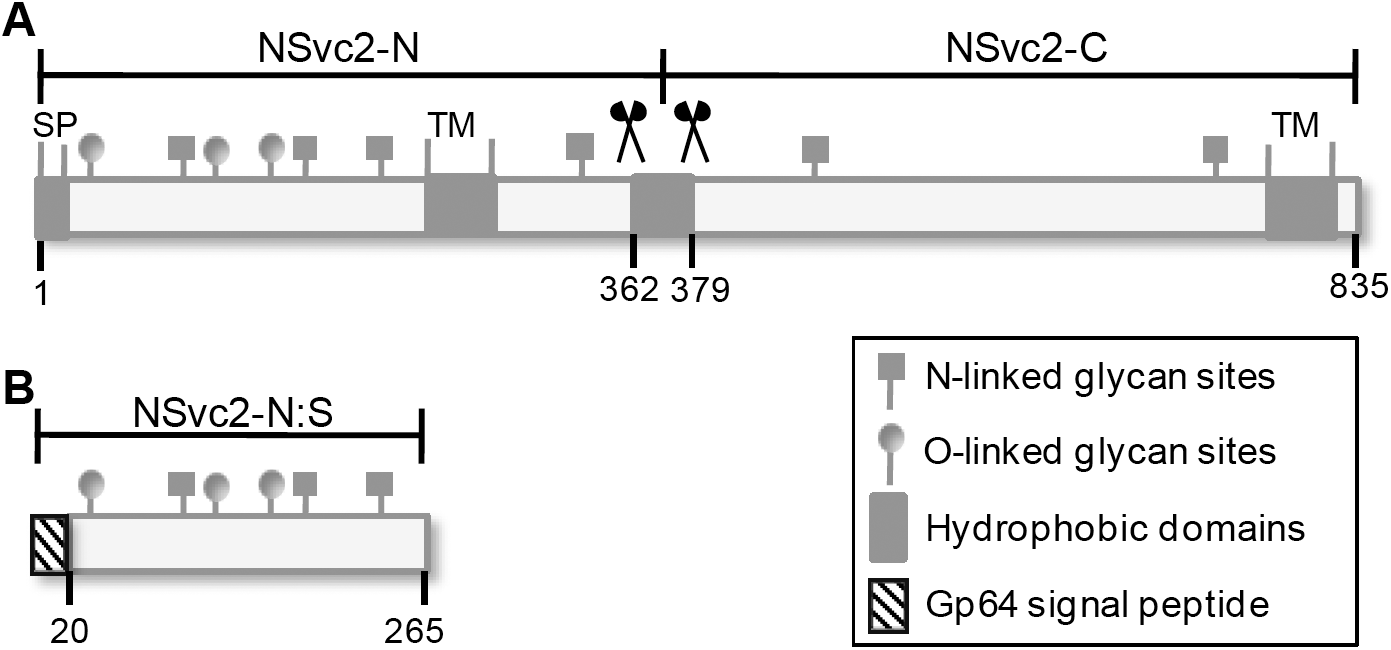
Schematic diagram showing the full length NSvc2 (A) and the recombinant soluble NSvc2-N (NSvc2-N:S) (B). Locations of the putative signal peptide, hydrophobic domains, and glycosylation sites are indicated. The signal peptide of NSvc2-N:S is replaced with the signal peptide of Gp64.

**Fig S2.**
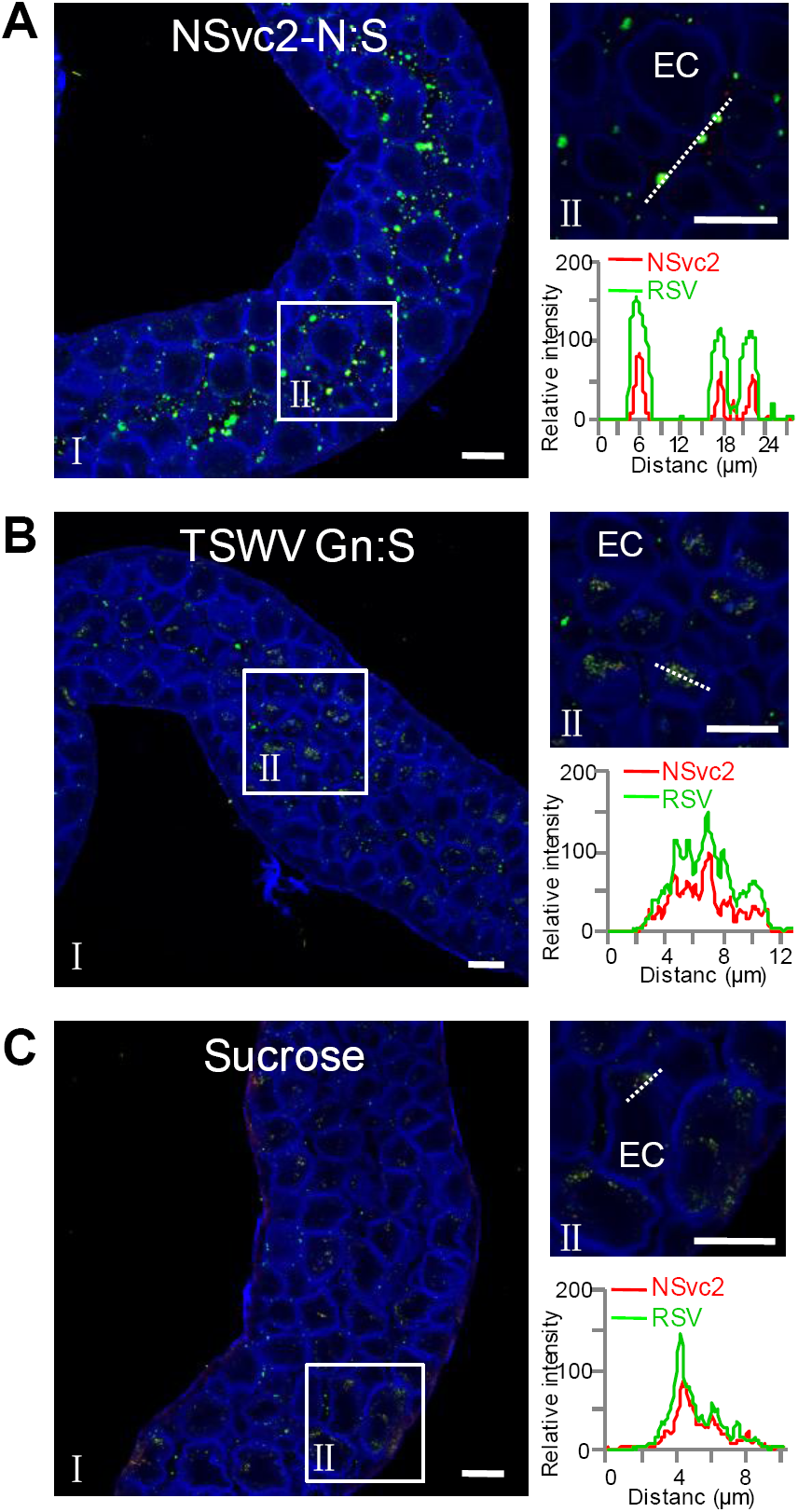
The effects of pre-incubation with purified NSvc2-N:S (A), TSWV Gn:S (B) and sucrose alone (C) on RSV entrance into SBPH midguts. The boxed regions are enlarged and shown on the right side. The overlapping fluorescence spectra were from the white dashed line indicated areas (right top and bottom panels). EC, epithelial cell; Bar, 25 μm.

**Fig S3.**
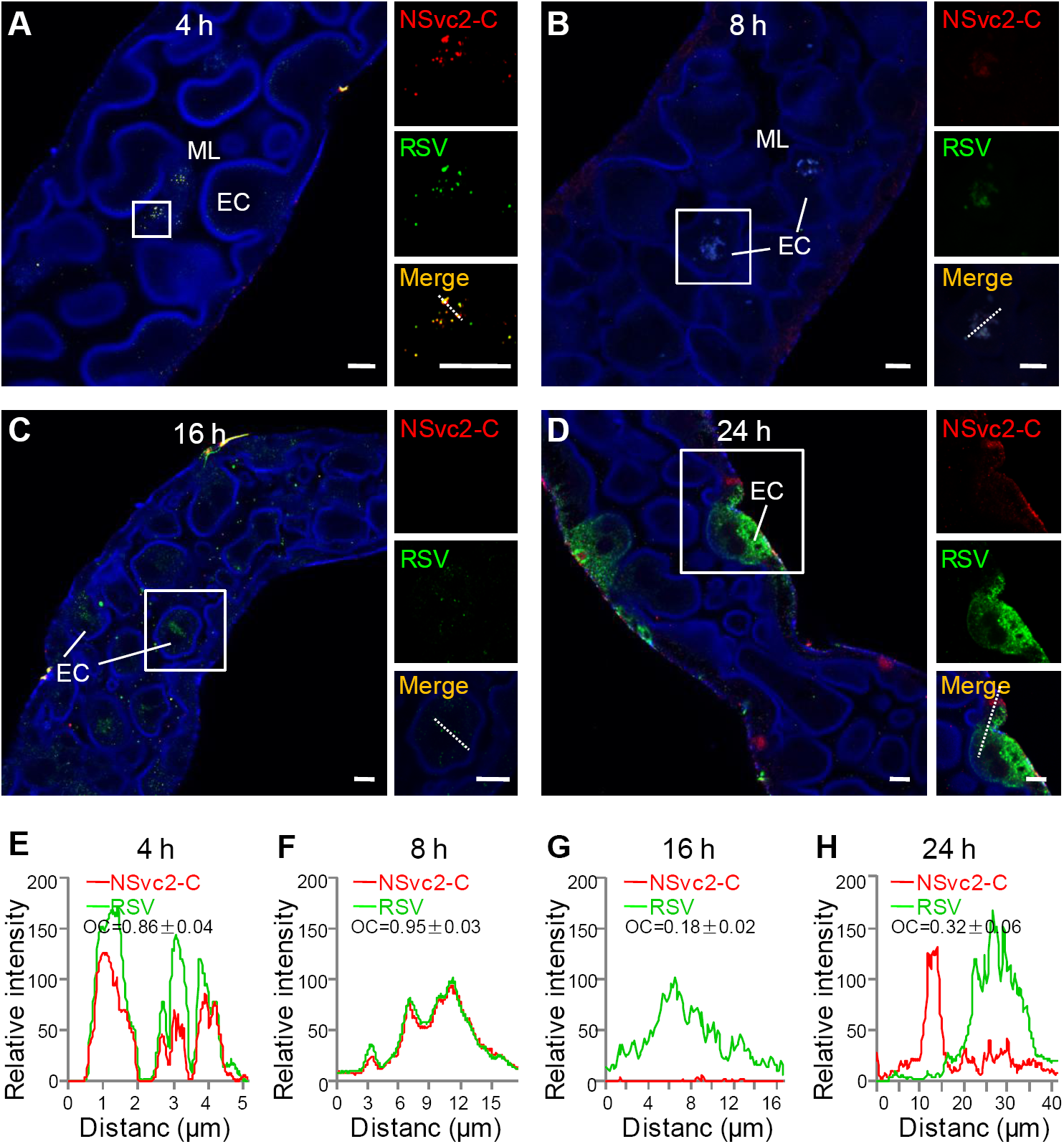
Time course study of NSvc2-C subcellular localization during RSV entrance into SBPH midgut. (A) NSvc2-C was co-localized with RSV virion on midgut microvillus surface (blue) at 4 h post feeding on RSV-infected seedlings. The boxed region was enlarged and shown on the right side. The labeled NSvc2-C is shown in red and the labeled RSV virion is shown in green. (B) NSvc2-C and RSV virion were co-localized in the endosomal-like vesicles in midgut epithelial cells at 8 h post infestation. (C) RSV virion was detected in cytoplasm of epithelial cells at 16 h post infestation but not NSvc2-C. (D) NSvc2-C was again not detected together with RSV virion at 24 h post infestation. (E–H) Analyses of overlapping fluorescence spectra from the white dashed lines indicated regions in the merged images. The overlap coefficient (OC) values were determined by the LAS X software. ML, midgut lumen; EC, epithelial cell; Bar, 10 μm.

**Fig S4.**
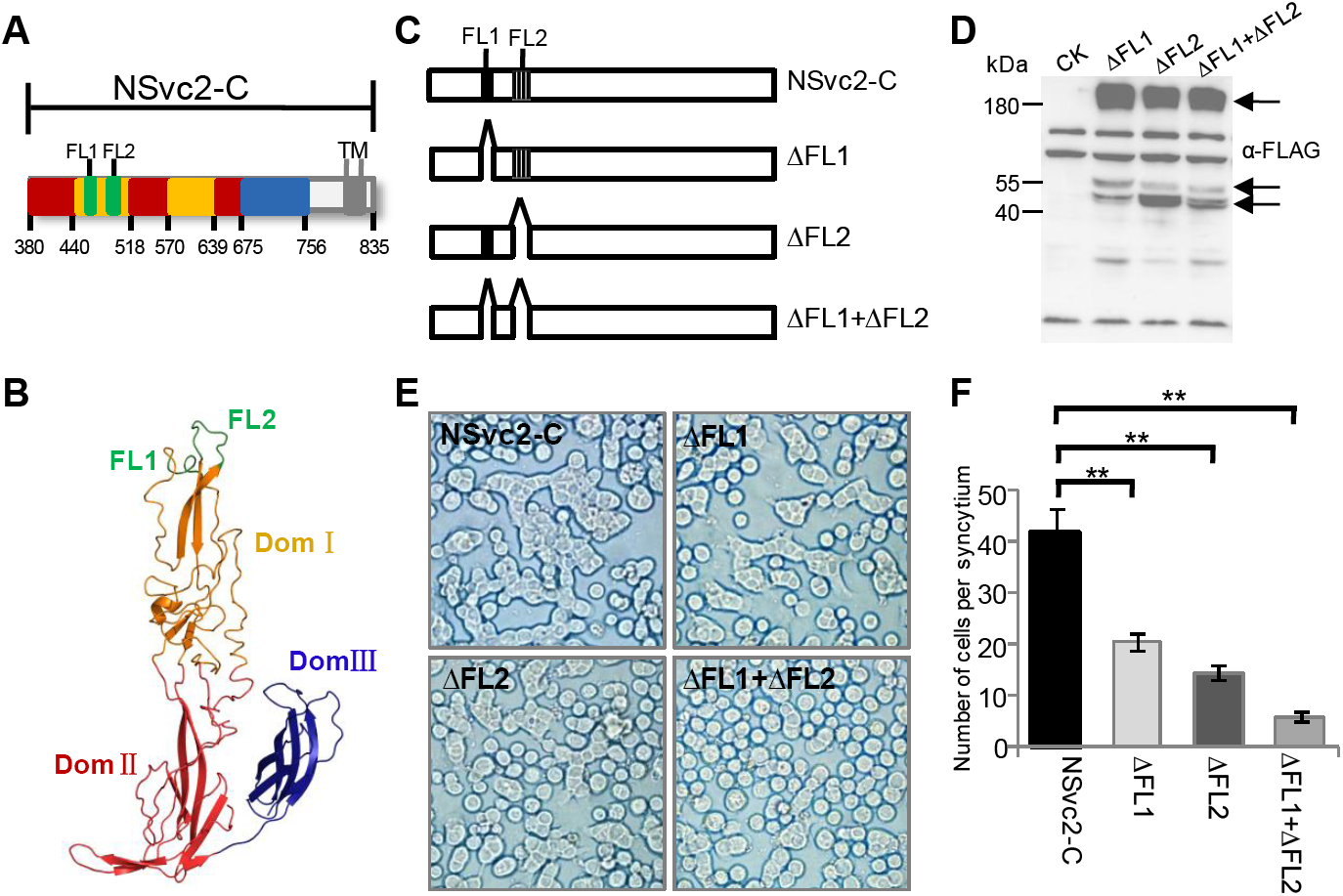
NSvc2-C hydrophobic fusion-loop motifs are required for cell-cell membrane fusion. (A) A diagram showing different NSvc2-C domains. Different domains are shown in yellow (domain I), red (domain II) and blue color (domain III). Fusion loops (FL1 and FL2) are shown in green, and a hydrophobic region in gray. (B) A three dimensional homology based model of NSvc2-C with the same color arrangement as shown in (A). (C) Schematic representations of the wild type and FL deletion NSvc2-C mutants. Locations of FL1 (black) and FL2 (gray) are shown. Deletions of one or double FLs are indicated with upward open arrows. (D) Expressions of FLAG-tagged WT or mutant NSvc2-C in Sf9 cells were determined by immunoblotting. The blots were probed with a FLAG-specific antibody. Arrows indicate the bands of the expressed NSvc2-C proteins. (E and F) Analyses of fusogenic activities of the WT or mutant NSvc2-C. Sf9 cells were infected with recombinant baculoviruses carrying the WT or deletion mutants. At 48 h post infection, the cells were treated with PBS, pH5.0, for the cell-cell membrane fusion assays (E). The numbers of cells showing syncytium were recorded and analyzed (F). The experiment was repeated three times. **, *p* < 0.01.

**Table S1.**
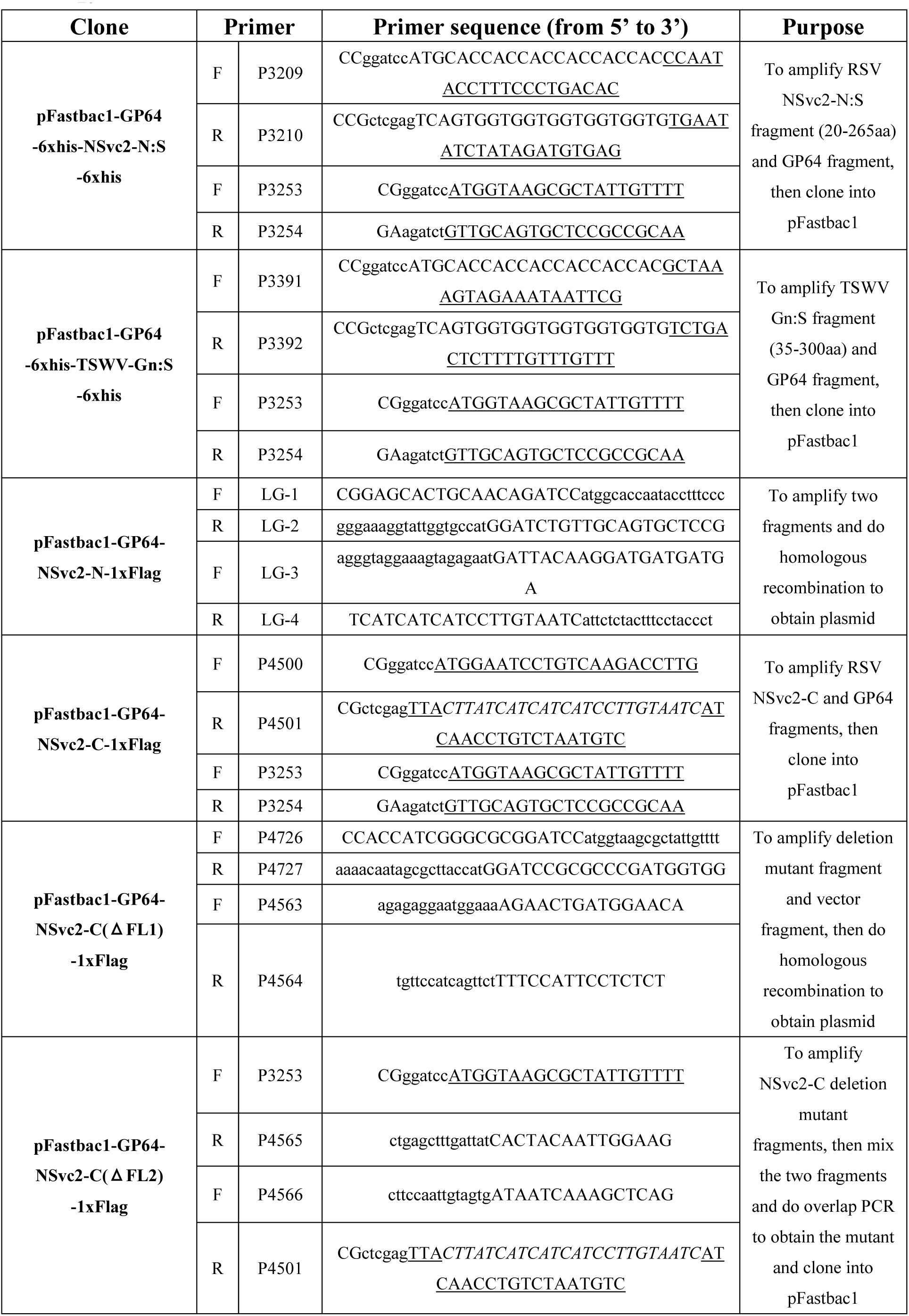
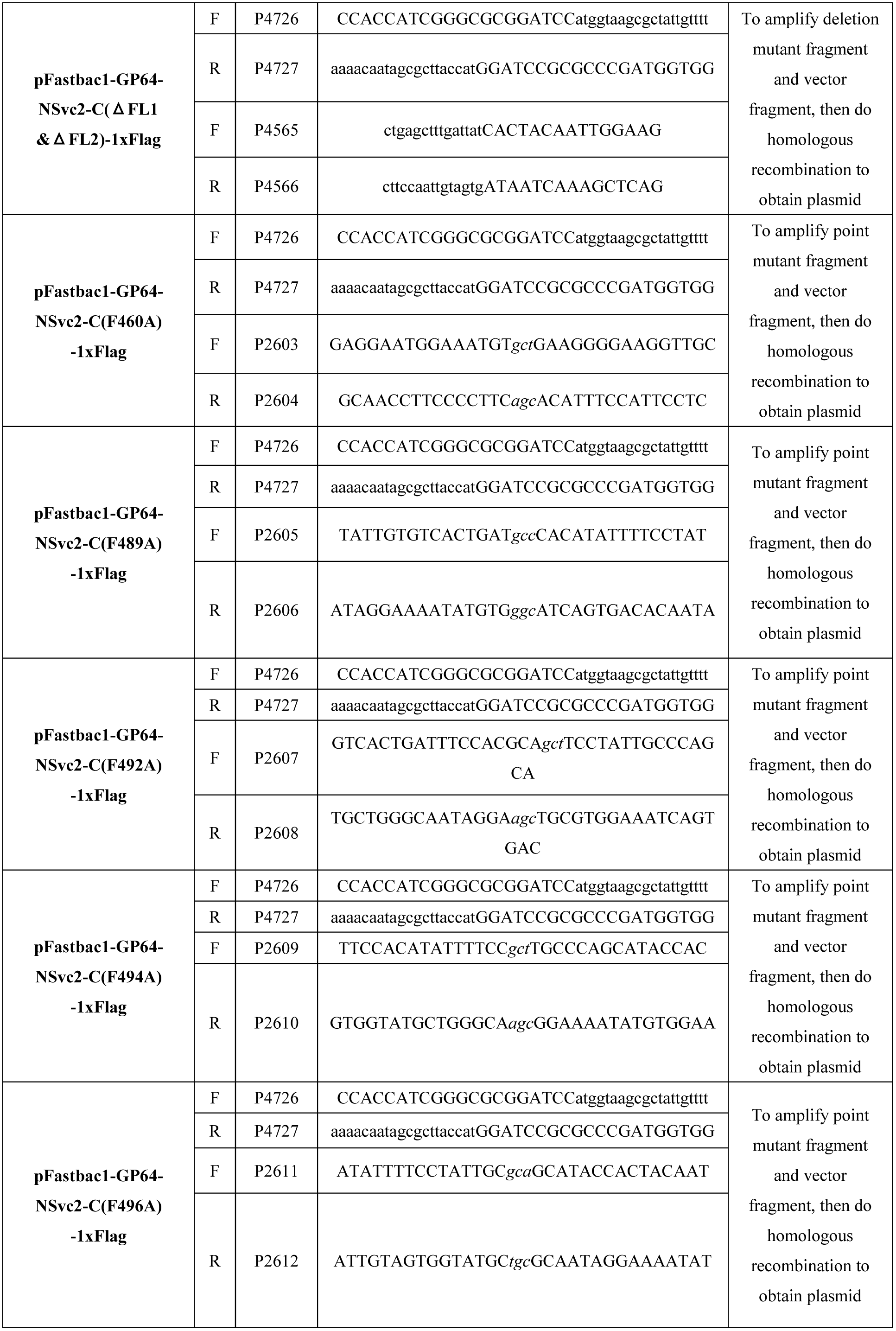
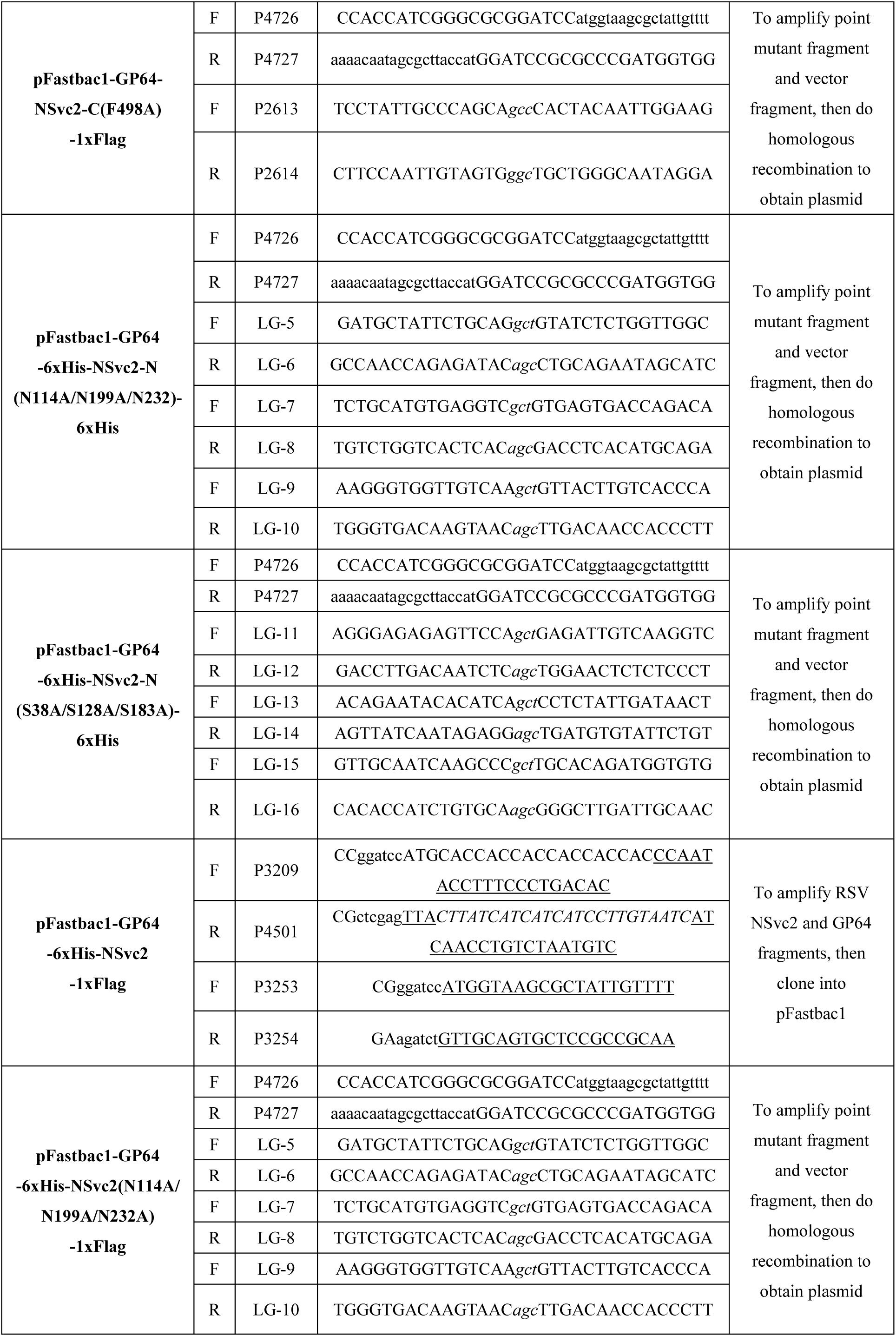
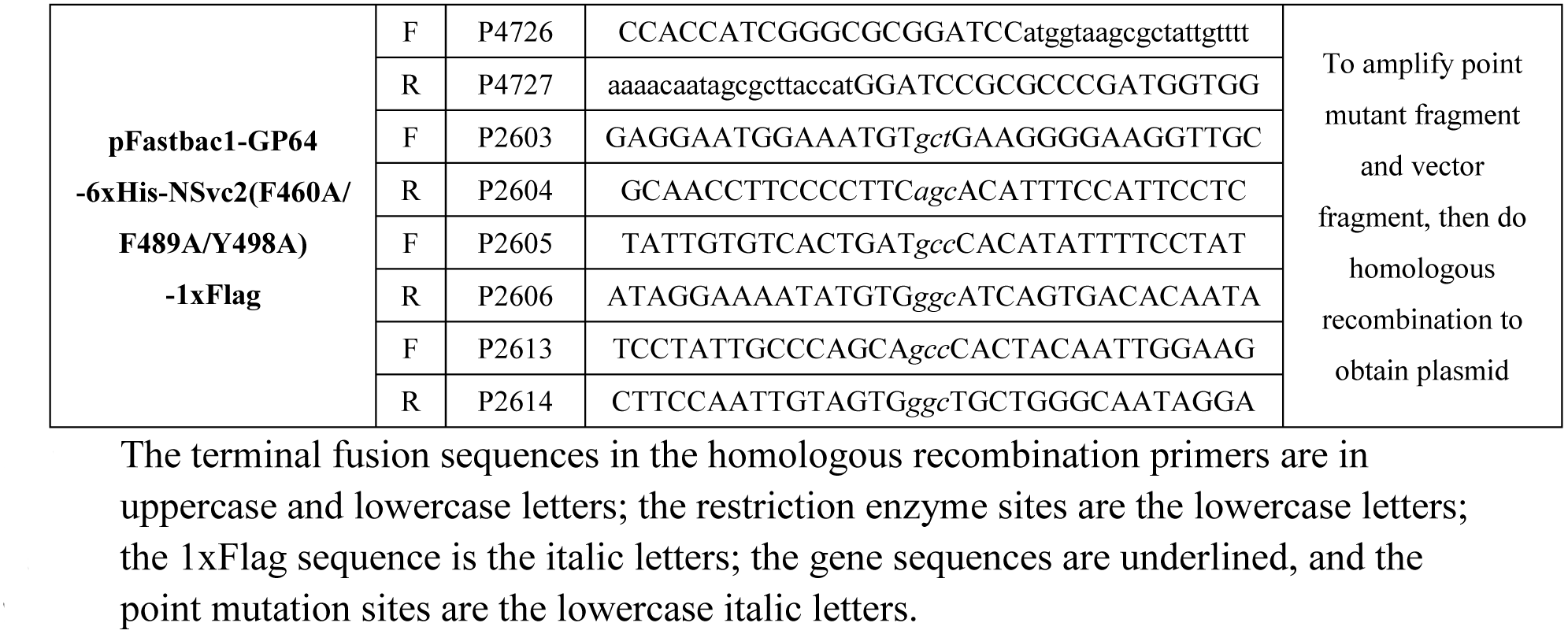
Primers used in this study

